# Bidirectional control of fear memories by the cerebellum through the ventrolateral periaqueductal grey

**DOI:** 10.1101/2020.02.19.956375

**Authors:** Jimena L. Frontera, Hind Baba Aissa, Romain William Sala, Caroline Mailhes-Hamon, Ioana Antoaneta Georgescu, Clément Léna, Daniela Popa

## Abstract

Fear conditioning is a form of associative learning that is known to involve brain areas, notably the amygdala, the prefrontal cortex and the periaqueductal grey (PAG). Here, we describe the functional role of pathways that link the cerebellum with the fear network. We found that the cerebellar fastigial nucleus (FN) sends glutamatergic projections to vlPAG that synapse onto glutamatergic and GABAergic vlPAG neurons. Chemogenetic and optogenetic manipulations revealed that the FN-vlPAG pathway controls bi-directionally the strength of the fear memory, indicating a role in the association of the conditioned and unconditioned stimuli, a function consistent with vlPAG encoding of fear prediction error. In addition, we found that a FN - thalamic parafascicular nucleus pathway, which may relay cerebellar influence to the amygdala, is involved in anxiety and fear expression but not in fear memory. Our results reveal the contributions to the emotional system of the cerebellum, which exerts a potent control on the strength of the fear memory through excitatory FN-vlPAG projections.

## Introduction

Fear conditioning is a form of associative learning in which an animal learns to associate the presence of a neutral stimulus (conditioned stimulus, CS) with the presence of an aversive fear eliciting stimulus (unconditioned stimulus, US). The amygdala complex, comprised of the basolateral complex (BLA), the central nucleus (CEA), and the intercalated cell masses (ITC), is one of the key brain regions for the fear memory acquisition and storage ^1–4^. Fear conditioning induces changes not only into the amygdala but also in auditory and multimodal nuclei of the thalamus, auditory cortex, medial prefrontal cortex (mPFC) and hippocampus ^2, 5–8^. In addition, fear extinction involves the formation of a new memory trace that attenuates fear responses to a conditioned aversive memory. The distributed network that controls fear extinction involves many of the brain areas that are important for fear conditioning ^2, 7, 9^.

The midbrain periaqueductal grey (PAG) is known to mediate both learned and innate freezing behavior via excitatory projections to the magnocellular medulla from the ventrolateral PAG (vlPAG)^10^. In learned freezing, the recruitment of these neurons is produced by a disinhibition triggered by GABAergic inputs from the amygdala in cued fear ^10^ and glutamatergic inputs from the mPFC in contextual fear ^6^. Some vlPAG projection neurons also participates to pain processing ^11–13^. However, vlPAG is not a simple regulator of sensory and motor functions, it is also involved in fear learning by generating a ‘fear prediction error’ assessing how unexpected is an incoming unconditioned stimulus^14^; this step is essential in implementing the Rescorla-Wagner rule which states that learning is driven by departure from expectation^15^.

Despite the well-described role of the cerebellum in motor coordination and sensorimotor integration, it is now well established that the cerebellum is also involved in a number of non-motor functions ^e.g. 16–19^. Notably, the cerebellum is strongly recruited in aversive emotional states: in humans, neuroimaging studies have revealed changes in cerebellar activation, also mainly in the vermis, in relation to negative emotions (e.g. during recall of self-generated emotional episodes ^20^). Consistent with this, localized cerebellar lesions, principally in the midline vermis, account in large part for the emotional disturbances, inappropriate behavior and changes in affect, which are collectively termed the “cerebellar cognitive affective syndrome” ^21^ and reported in cerebellar patients. Several studies have shown that the cerebellum has functional connections from fear-related areas including the PAG, the amygdala, and the prefrontal cortex ^22–25^. In accordance with the existence of such connections, Pavlovian fear conditioning affects cerebellar plasticity ^26^, post-conditioning cerebellar inactivation affects memory consolidation^27^, and cerebellar lesions -or inactivation-modulate freezing ^24, 28;^ however, the pathways by which the cerebellum participates to fear learning or expression remain undefined.

The cerebellar vermis, which is most consistently associated with emotional pathologies and fear expression ^20, 24, 29^, projects to the fastigial nucleus (FN), a deep cerebellar nucleus (DCN) which projects to many targets from the spinal cord to the diencephalon^22^. The purpose of the present study is to study the contribution of specific FN output pathways to fear learning. Using neuroanatomical tracings, chemogenetic modulation of the cerebellar input to the vlPAG and to the BLA during fear conditioning, optogenetics and extracellular electrophysiological recordings in awake freely moving animals, we demonstrate the contribution of the cerebellum to fear learning through its inputs to the vlPAG.

## Results

### Neuroanatomical link between the cerebellum and vlPAG

In order to examine cerebellar projections to areas involved in the fear circuitry, we used anterograde viral tracing and we have identified possible target areas of the projections from the FN of the cerebellum. We stereotaxicaly infused AAV-mCherry in the FN and we found projections exhibiting varicosities into the vlPAG (Fig.1 a-f), which receives inputs from the CEA to drive freezing behavior ^10^. Retrograde tracing from vlPAG also indicated the existence of monosynaptic projections from the FN, as well as from the CEA, from the PL and IL areas of the mPFC (Supplementary Fig. 1 a-e). Quantification of retrograde labeling from the vlPAG showed that at least 11.2 ± 0.8 % of neurons in the FN projects to vlPAG (n=4; Fig. 1 g-i), suggesting that this pathway plays an important contribution to cerebellar control of the emotional system ^22^. Moreover, combining a *vglut2-cre* mice line with the infusion of AAV-DIO-tdTom in FN and retrograde AAV-GFP in vlPAG allowed us to determine that the FN-vlPAG projecting neurons correspond to vesicular glutamate transporter 2 expressing (vGluT2+) neurons (85.4 ± 4.0 % of GFP-expressing cells co-localized with vGluT2+ neurons, n=4; Fig. 1 j-l). On the other hand, the retrograde tracing in *glyt2-gfp* mice failed to reveal the FN–vlPAG glycinergic projecting neurons (none of retrograde stained cell co-localized with GlyT2+ neurons in the FN, n=3; Supplementary Fig. 1f, g). Moreover, FN GABAergic neurons are known to only project to the inferior olive ^30^. Overall, these results indicate that the vlPAG receives glutamatergic monosynaptic inputs from the FN of the cerebellum.

**Figure 1.**
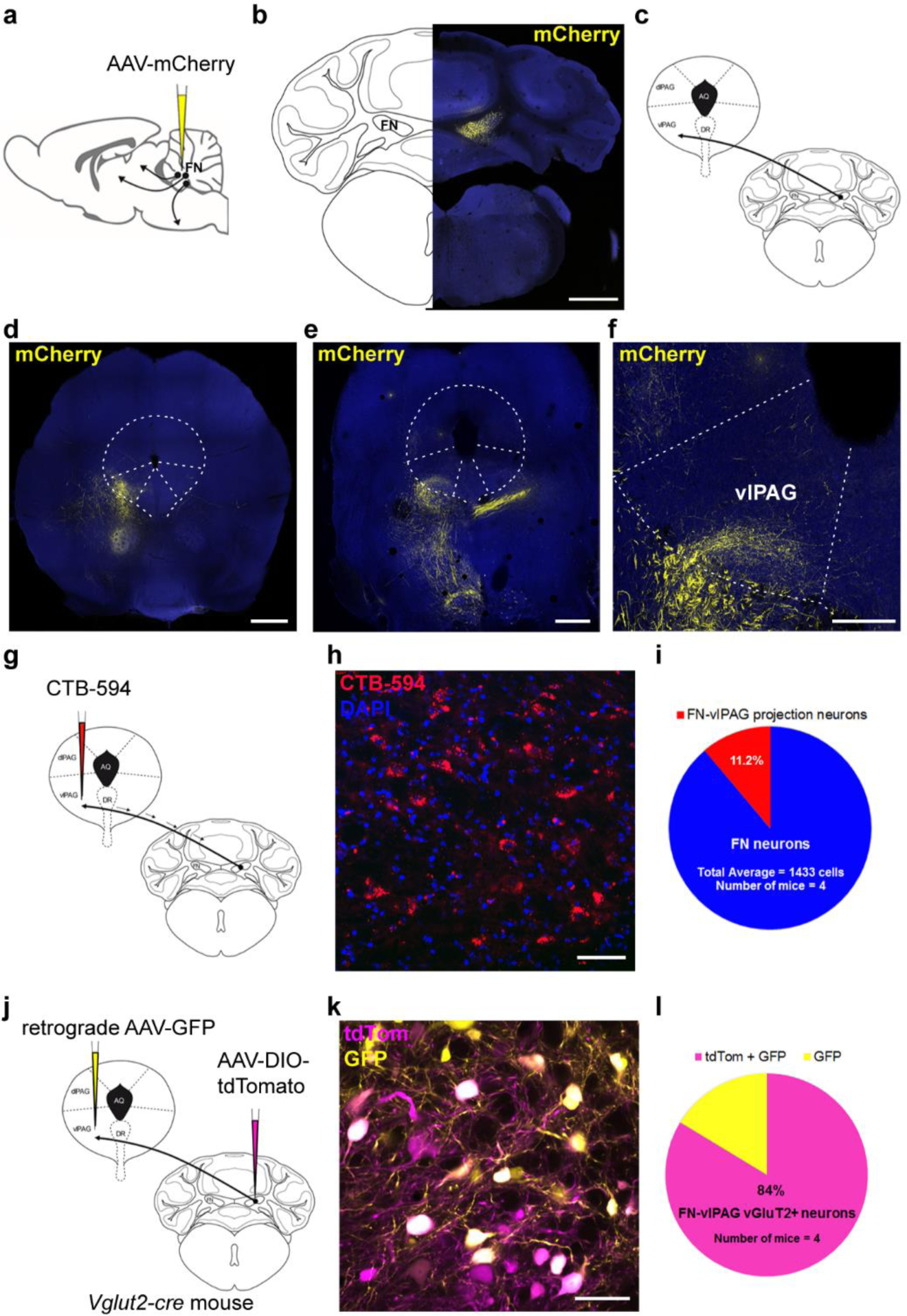
Cerebellum sends excitatory monosynaptic projections to vlPAG. **a**, Anterograde tracing strategy by AAV-mCherry injection in the FN. **b**, Expression of anterograde AAV-mCherry in the FN of the cerebellum (scale bar, 1mm).**c**, Schematic representation of anterograde tracing of AAV-mCherry from the FN into vlPAG. **d-e**, Anterograde expression of mCherry in FN neuronal projections exhibiting varicosities in the vlPAG (scale bar, 500 µm), anterior (d) and medial (e) PAG sections. **f**, Zoom-in from d (scale bar, 250 µm). **g**, Retrograde labeling approach by injecting CTB-594 in the vlPAG. **h**, Retrograde labeling of FN-vlPAG projecting neurons with CTB-594, co-staining with DAPI (scale bar, 50 µm). **i**, Quantification of retrograde labeled FN neurons projecting to the vlPAG (n=4 mice, total average= 1433 cells). **j**, Viral injection of retrograde AAV-GFP and anterograde AAV-DIO-tdTom into *vglut2-cre* mice strategy. **k**, Co-expression of anterograde AAV-DIO-tdTom and retrograde AAV-GFP in vGluT2+ FN neurons projecting to vlPAG (scale bar, 50 µm). **l,** Quantification of co-localizing retrograde AAV-GFP and AAV-DIO-tdTomato vGluT2+ neurons in the FN.

### Glutamatergic FN input onto glutamatergic and GABAergic vlPAG neurons

To determine whether the inputs from the FN synapse either onto excitatory or inhibitory neurons in the vlPAG, we analyzed the axonal projections and synaptic boutons of FN vGluT2+ neurons. We localized glutamatergic FN terminals by cre-dependent viral expression of the presynaptic marker synaptophysin-GFP in adult *vglut2-cre* mice (Fig. 2 a, e). In addition, we identified glutamatergic neurons in the vlPAG by infusion of AAV-DIO-tdTomato, or the GABAergic neurons by immunostaining of Gad67. Glutamatergic FN terminals were found to contact both glutamatergic and Gad67+ neurons in vlPAG (Fig. 2 a-d and e-h, respectively). Therefore, these results indicate that FN-vlPAG glutamatergic projections target both glutamatergic and GABAergic vlPAG neurons.

**Figure 2.**
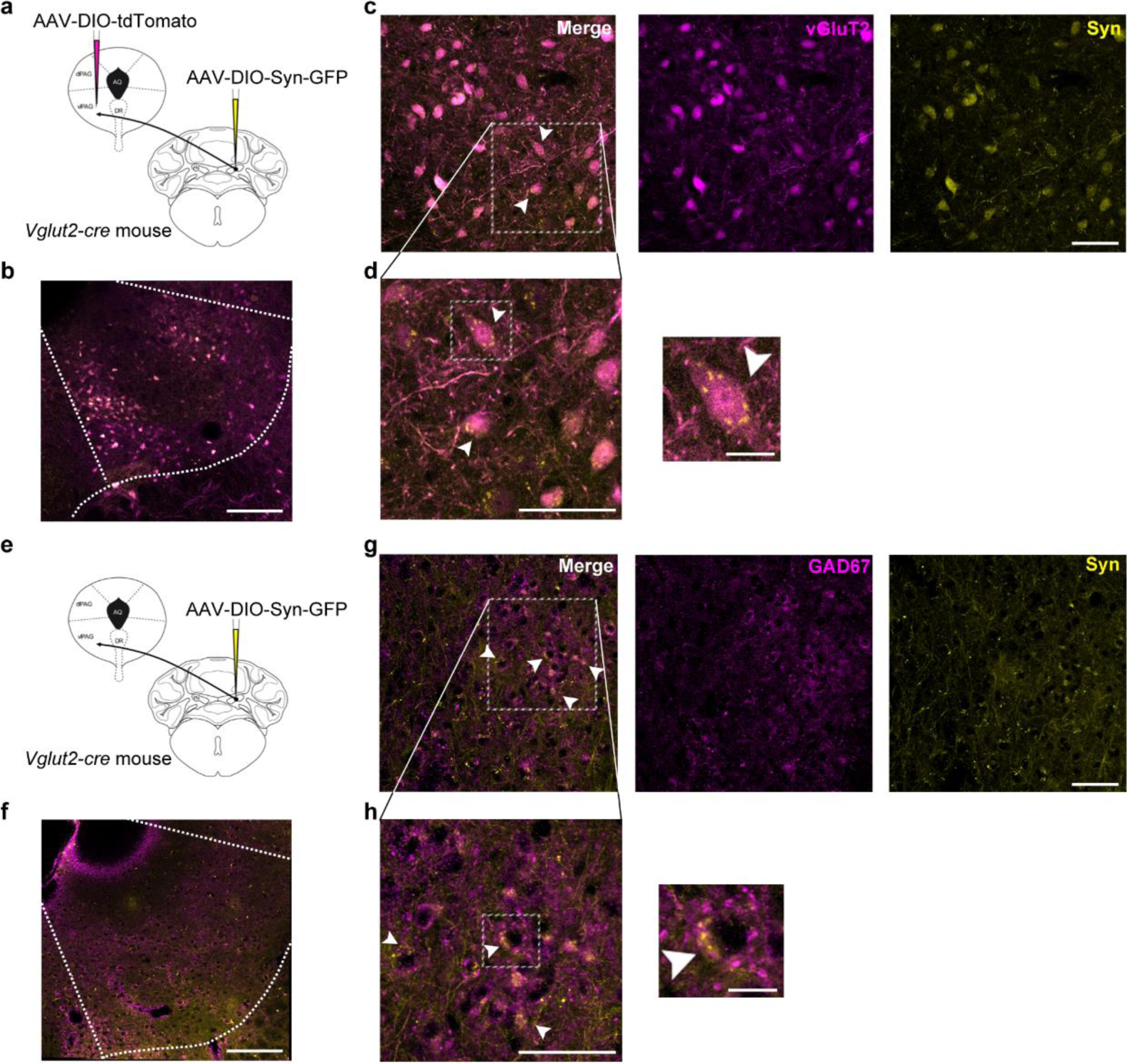
Glutamatergic FN input onto excitatory and inhibitory neurons in the vlPAG. **a**, Injection of cre-dependent AAV-DIO-Syn-GFP and AAV-DIO-tdTomato in vglut2-cre mice line strategy. **b,** Expression of tdTomato in vGluT2+ neurons and Syn-GFP in FN terminals in the vlPAG (scale bar, 250 µm). **c,** FN glutamatergic synapses contacting vGlut2+ neurons in vlPAG identified by expression of Syn-GFP (arrowheads, scale bar, 50 µm). Some of the somatic labeling likely reflects the high somatic tdTomato expression and crosstalk detection with the GFP; intense Synapsin synaptic labeling is best visualized in high magnification panel. **d,** High-magnification from c (left panel, scale bar, 50 µm) and zoom-in of box area (right panel, scale bar, 12.5 µm). **e,** Injection of cre-dependent AAV-DIO-Syn-GFP in vglut2-cre mice line strategy. **f,** Immunostaining of GAD67 and Syn-GFP expression in FN terminals in the vlPAG (scale bar, 250 µm). **g,**FN glutamatergic projections synapsing on vlPAG GABAergic neurons identified by co-staining of Syn-GFP and GAD67 in vlPAG (arrowheads, scale bar, 50 µm). **h,** High-magnification from g (left panel, scale bar, 50 µm) and zoom-in of box area (right panel, scale bar, 12.5 µm).

### Optogenetic stimulation of FN neurons induces short latency response in vlPAG cells

To study the impact of optogenetic FN stimulation in the vlPAG we expressed channel-rhodopsin-2 (ChR2) specifically in FN-vlPAG projecting neurons by local injection of cre-dependent AAV-DIO-ChR2-GFP in the FN, combined with local injection of retrograde AAV-cre-EBFP into the vlPAG (Supplementary Fig. 2 a, b). Then, we stimulated FN-vlPAG neurons via FN illumination and we recorded vlPAG activity in awake freely moving animals (Fig. 3 a-f). Under FN stimulation, we found LFP negative deflection (Fig. 3 a) and cells exhibiting an increase of firing in vlPAG contra-lateral to the FN stimulation (Fig. 3 b,c) where the FN is preferentially sending projections. The ramp of stimulation exhibited a range of vlPAG responses, which increased as a function of light intensity (Fig. 3 d). Responses at low intensities were found in the vlPAG contra-lateral to the stimulated FN, while increasing intensities recruited cells on the ipsi-lateral vlPAG (Fig. 3 e). The latencies of response at the highest intensity range from 7 to 50 ms (Fig. 3 f), suggesting the existence of direct and indirect excitation of vlPAG by FN input. These results demonstrate the existence of a functional connectivity between FN and the vlPAG.

**Figure 3.**
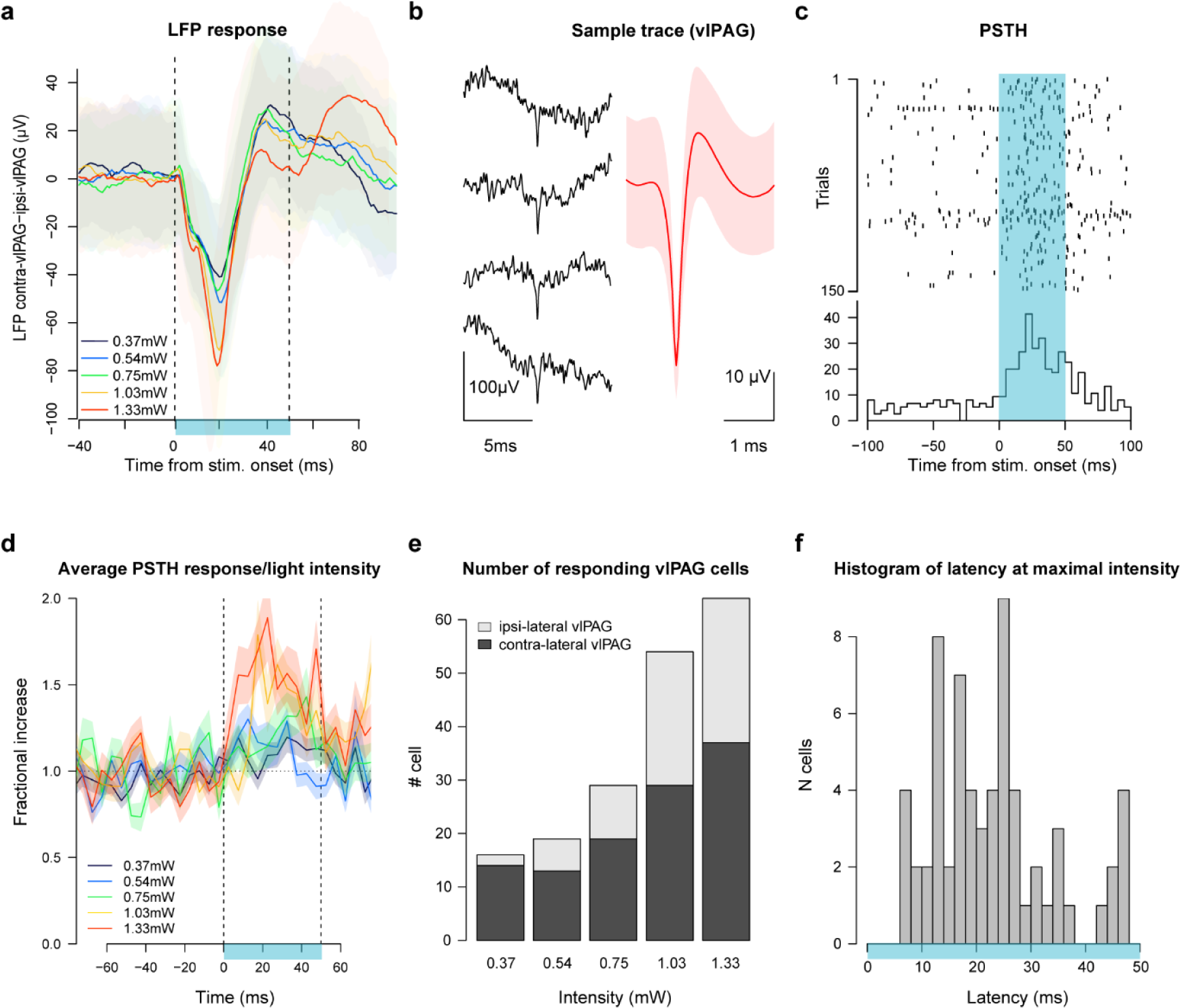
Optogenetic FN stimulation induces short latency responses in vlPAG. **a**, LFP response in vlPAG to FN stimulation for 5 different light intensity showed with different colors. **b,** Excerpt of traces and spikes from vlPAG; right: average waveform for the neuron sorted on the channel; shaded area represent the standard deviation of the trace. **c,** Raster plot for a cell recorded in vlPAG around the 50 ms period of FN stimulation. Blue area indicates periods with cerebellar optogenetic stimulation. **d,** Light-dependent effect for the cells in the vlPAG that increased their firing during 50 ms light stimuli at each intensity bin=5 ms (n=61 out of 222 neurons from 4 mice with significant response at the highest intensity; significance is established with 2 ms bin, threshold used to identify significance in PSTHs is: 4.1xSd.). **e,** Number of responding vlPAG cells (ipsi and contra-lateral) to FN stimulation. **f,** Latencies of responses in cells in vlPAG at highest intensity during the 50ms of FN stimulation.

### Neuroanatomical link between the cerebellum and the amygdala

The BLA is a key brain structure for fear memory acquisition and consolidation. Therefore, in order to evaluate a possible neuroanatomical connection between the cerebellum and the BLA we combined anterograde tracing from the FN with retrograde tracing from the BLA (Fig. 4 a-c). This analysis failed to reveal the presence of BLA projecting neurons in the vlPAG surrounding the area where the FN fibers were localized. However, neuronal projections from FN were found in the parafascicular nucleus of the thalamus (PF) together with BLA projecting neurons (Fig. 4 d, e), suggesting that PF might be a relay area between the cerebellum and the BLA.

**Figure 4.**
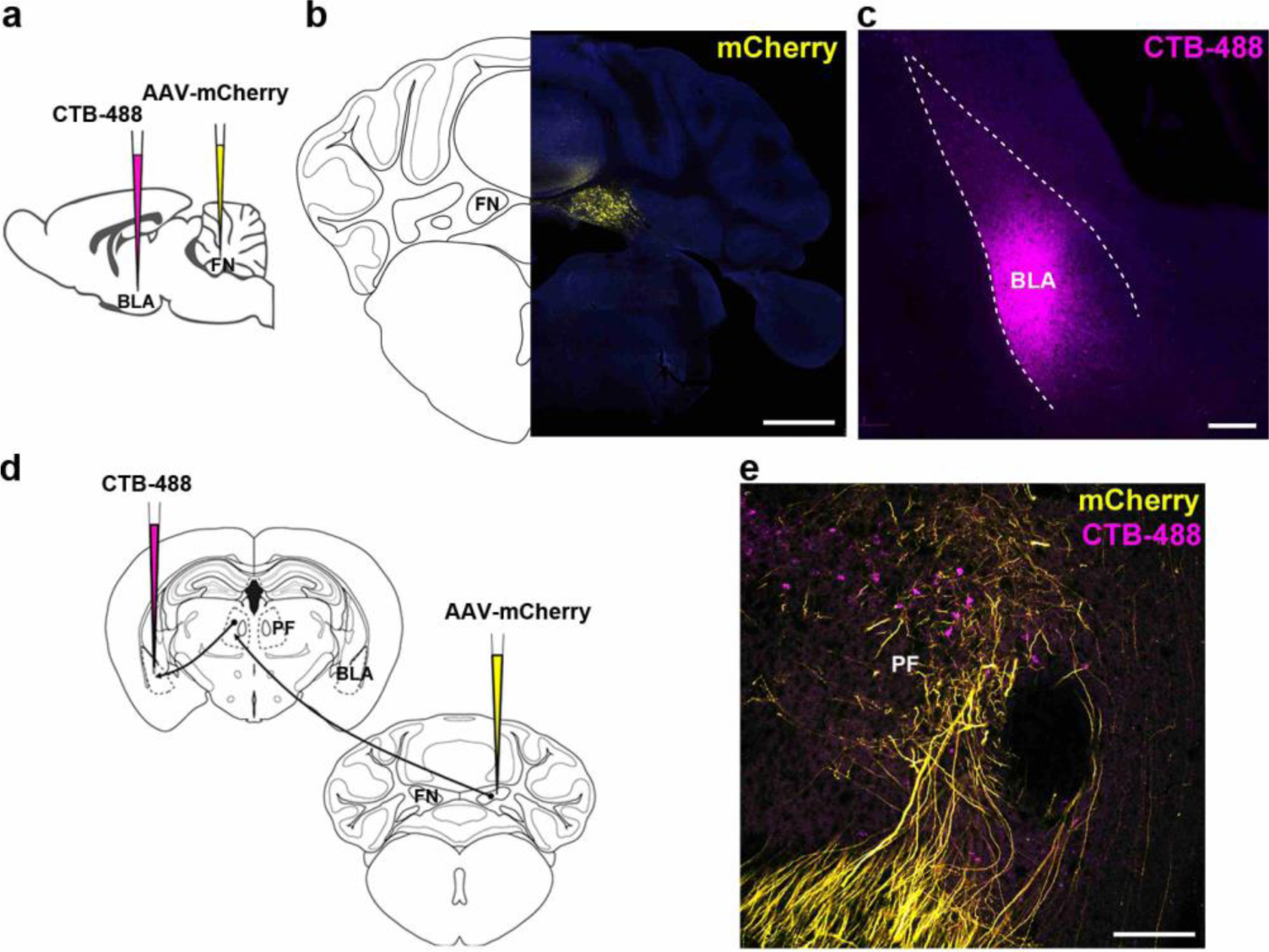
Cerebellar link to the BLA through thalamic PF. **a**, BLA injection of retrograde CTB-488 in combination with anterograde AAV-mCherry in the FN strategy. **b**, Expression of anterograde AAV-mCherry in the FN (scale bar, 1mm). **c**, Infusion of CTB-488 in the BLA (scale bar, 200 µm). **d**, Retrograde tracing from BLA combined with anterograde tracing from the FN converged at the thalamic PF nucleus. **e**, PF neurons projecting to the BLA surrounded by fibers from FN (scale bar, 200 µm).

### Cerebellar output to vlPAG controls fear memory formation

To determine whether the FN-vlPAG and FN-PF pathway plays a role in Pavlovian fear conditioning, we examined the effect of activating or inhibiting transiently these projections. First, we expressed excitatory or inhibitory DREADD receptors activated by CNO in neurons that target the vlPAG, by infusing cre-dependent anterograde AAV-hM3Dq or -hM4Di in FN in combination with the infusion of retrograde CAV2-cre in the vlPAG (Fig. 5 a-c). We performed a classical Pavlovian fear conditioning protocol, which consisted in 5 CS-US presentations, followed by three extinctions sessions of 25 CS, and a final recall test of 5 CS presentations one week after to evaluate long-term maintenance of fear and extinction memories (Fig. 5 d). Mice injected with saline and expressing the excitatory or the inhibitory DREADDs had similar freezing levels (Supplementary Fig. 3 a), thus they were grouped together in the control (injected with saline) group. In addition, we checked that CNO had no effect *per se* on the freezing behavior by performing the fear and extinction protocols on control sham mice. Mice exposed to CNO indeed exhibited the same freezing levels than the control mice, which received saline (Supplementary Fig. 3 b). Moreover, chemogenetic activation or inhibition of FN-vlPAG pathway had not effect on the immobility state of the mice in an open field compared to the saline group (Supplementary fig. 3 c), neither on the freezing levels in the context A before fear conditioning (Supplementary fig. 3 d) or in the context B before extinction training (Supplementary fig. 3 e). These results indicate that FN-vlPAG pathway does not affect motor responses as immobility state or freezing behavior. Therefore, we first analyzed the role of these connections during fear conditioning. For this purpose, we activated or inhibited FN-vlPAG projecting neurons with CNO during the acquisition phase. Under this condition, only the inhibition of FN-vlPAG projections had significant effect on fear response, by increasing mildly the freezing at the end of the fear conditioning (Fig. 5 e), while mice under FN-vlPAG pathway activation exhibited similar fear response than the control group. Strikingly, during the first extinction training the following day, mice in which the FN-vlPAG pathway had been activated during fear conditioning, exhibited a drastic drop in freezing behavior along the CS presentations, reaching basal freezing levels already at the end of the first extinction session (Fig. 5 e). During the second extinction training, this group of mice succeeded to retrieve the fear memory at the first CS presentation but they reached basal levels of freezing at the third CS presentation, indicating a faster extinction than in the control mice. In contrast, mice in which the FN-vlPAG pathway had been inhibited presented a stronger freezing behavior throughout the first extinction training as well as for the second and third extinction sessions, compared to the control mice (Fig. 5 e). Moreover, when we evaluated the long-term maintenance of the fear and extinction memories by a recall session of five CS presentations one week after, mice that underwent

**Figure 5.**
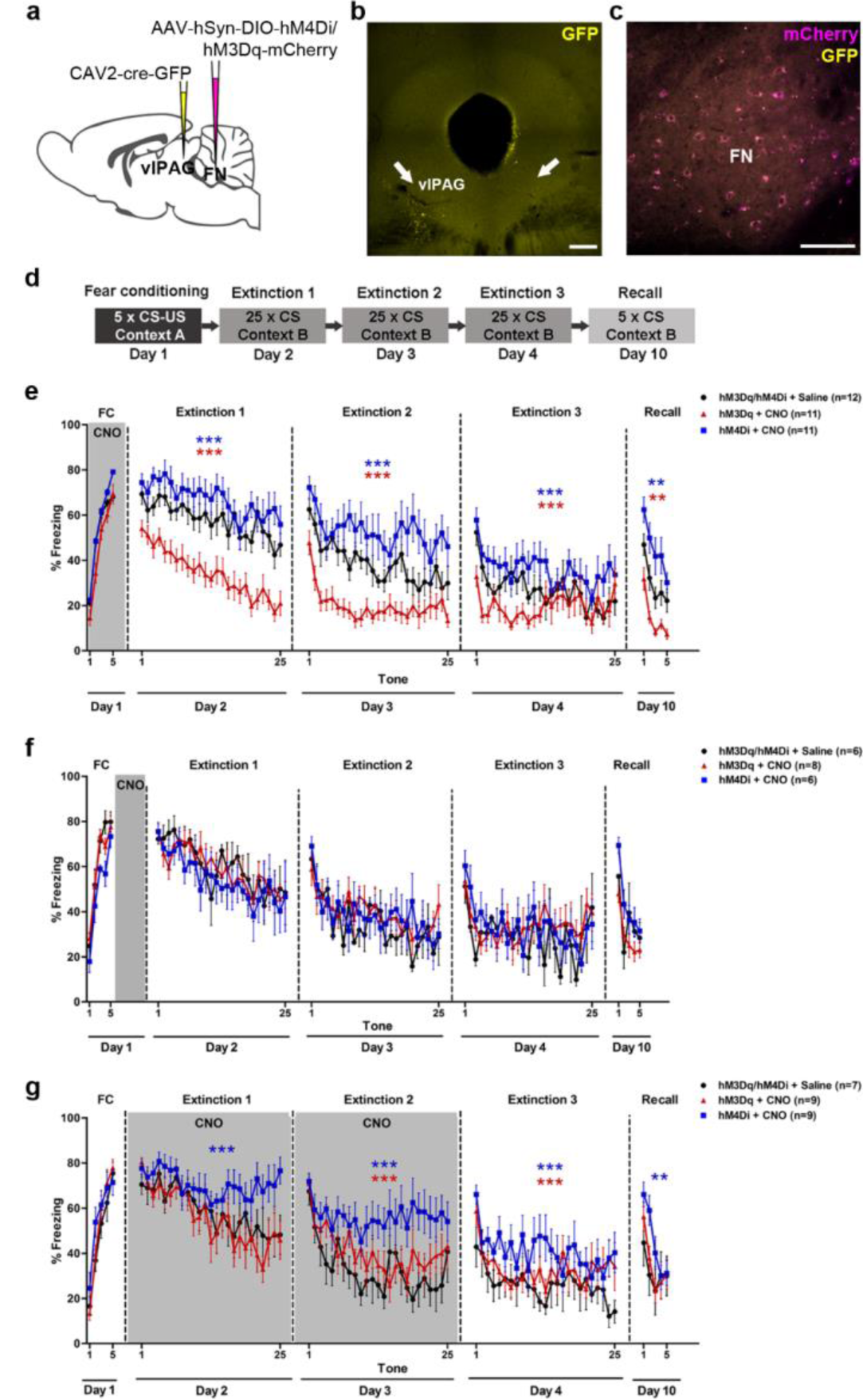
FN-vlPAG projections control fear memory formation and extinction learning. **a**, Chemogenetic strategy to modulate the activity of FN-vlPAG projecting neurons, bilateral cre-dependent expression of excitatory or inhibitory DREADDs (hM3Dq or hM4Di), respectively) in the FN in combination of retrograde CAV2-cre-GFP injection in the vlPAG. **b**, Injection site of CAV2-cre-GFP in the vlPAG (arrows, scale bar, 200 µm). **c**, Cre-dependent expression of DREADDs in the FN (scale bar, 100 µm). **d**, Classical fear conditioning and extinction protocol used. **e**, FN-vlPAG pathway stimulation (hM3Dq + CNO, n = 11) or inhibition (hM4Di + CNO, n = 11) during fear conditioning. Stimulation of FN-vlPAG pathway during fear conditioning provoked a decrease in freezing levels during the extinction sessions (Extinction 1: F(1, 546) = 251.5, Extinction 2: F(1, 546) = 159.3, Extinction 3: F(1, 546) = 25.9, P < 0.001, two-way ANOVA), while inhibition of this pathway induced an increase of freezing levels at the end of the fear conditioning (P < 0.05, Bonferroni post-hoc test) and along the extinction sessions (Extinction 1: F(1, 546) = 27.5; Extinction 2: F(1, 520) = 46.5, Extinction 3: F(1, 546) = 25.3, P < 0.001, two-way ANOVA), compared to the control group (hM3Dq/ hM4Di + Saline, n = 12). During the recall of fear and extinction memories, stimulated mice expressed less fear than the control group (Recall: F(1, 120) = 0.36, P < 0.001, two-way ANOVA), while inhibited mice expressed higher levels of fear expression (Recall: F(1, 102) = 16.1, P < 0.001, two-way ANOVA). **f**, Stimulation and inhibition of FN-vlPAG pathway during consolidation phase post-fear conditioning had not effect on freezing behavior (hM3Dq/ hM4Di + Saline, n = 6; hM3Dq + CNO, n = 8; hM4Di + CNO, n = 6; two-way ANOVA). **g**, Stimulation and inhibition of FN-vlPAG pathway during first and second extinction sessions. Inhibition of FN-vlPAG pathway induced higher levels of freezing behavior during the extinction sessions under effect of CNO and along the third extinction session and recall test without CNO, compared to the control group (hM3Dq/ hM4Di + Saline, n = 7; hM4Di + CNO, n = 9; Extinction 1: F(1, 364) = 37.9; Extinction 2: F(1, 364) = 96.9, Extinction 3: F(1, 364) = 44.2, P < 0.001; Recall: F(1, 78) = 10.5, P < 0.01, two-way ANOVA). Stimulation of this pathway during the extinctions also induced an increase of freezing behavior during the second and third extinction sessions (hM3Dq/ hM4Di + Saline, n = 7; hM3Dq + CNO, n = 9; Extinction 2: F(1, 364) = 12.7, Extinction 3: F(1, 364) =12.0, P < 0.001, two-way ANOVA). Lines represent means ± s.e.m. #P < 0.05 Bonferroni post-hoc test; **P < 0.01***P < 0.001, two-way ANOVA.

FN-vlPAG activation during fear conditioning exhibited a lower recall of fear memory than the control mice, while mice that underwent FN-vlPAG inhibition exhibited an increased fear memory compared to the control mice. These results suggest that FN-vlPAG projections contribute to the formation of conditioned fear memories, in which the activation of this pathway during the fear conditioning would decrease the strength of the fear memory formation, while the inhibition of this pathway would increase it. Consequently, the extinction of fear response during the extinction session is faster or slower, respectively.

In addition, we found an increase of c-Fos+ vlPAG neurons following fear conditioning when FN-vlPAG pathway was chemogenetically activated, compared to the control (Supplementary Fig. 4 a,b), confirming that FN-vlPAG glutamatergic projections induce an increase in neuronal activity when it is stimulated during fear conditioning.

Since the systemic CNO administration exerts an effect on neuronal activity for hours ^31–33^, we wondered whether the bidirectional control of FN-vlPAG pathway on fear memory occurs specifically during the acquisition or during the consolidation phase that immediately followed ^34, 35^. To answer this question we compared the effect of activating and inhibiting the FN-vlPAG pathway starting from the acquisition phase or from the consolidation phase. Then, we evaluated the fear response during the following recall and extinction sessions. We surprisingly found that activating and inhibiting the FN-vlPAG pathway only during the consolidation phase of fear memory had no effect on the strength of the fear memory (Fig. 5 f). Mice with FN-vlPAG activation or inhibition exhibited similar levels of fear response than the control group, during the recall and along the extinctions sessions. Taken together, these results indicate that the bidirectional control of the FN-vlPAG pathway of the fear memory occurs during the acquisition phase of the fear memory, in which the association of the CS-US is taking place.

Therefore, to assess the neuronal correlates of fear learning in FN and vlPAG, we recorded cellular activity during the extinction of the conditioned fear response (Fig. 6 a-c). We recorded 339 units in these structures during the 3 days of extinction learning (FN: n=181, vlPAG: n=158 from 6 hM3Dq + saline- and 7 hM3Dq + CNO-treated mice, Supplementary fig. 5 a). In the vlPAG, several classes of excitatory/inhibitory responses to the CS were found, in accordance with previous observations ^10, 12, 36;^ these responses were clustered (see Methods) according to their response profiles to the CS (Fig. 6, Supplementary fig. 5 b-d), yielding groups with 1) a fast excitatory response to the CS onset with progressive return to baseline, 2) a lasting inhibitory response to CS, 3) a late or sustained excitatory response to the CS, 4) a group with little if any modulation by the CS. Similar classes of responses in the FN could be revealed by grouping the PSTHs using the centroids of the clusters of the vlPAG (Fig. 6 e, f). The FN thus hosts different sets of cells that show parallel sensitivity to the CS to the ones observed in the vlPAG, indicating the existence of parallel firing time evolution of subpopulations in the vlPAG and FN.

**Figure 6.**
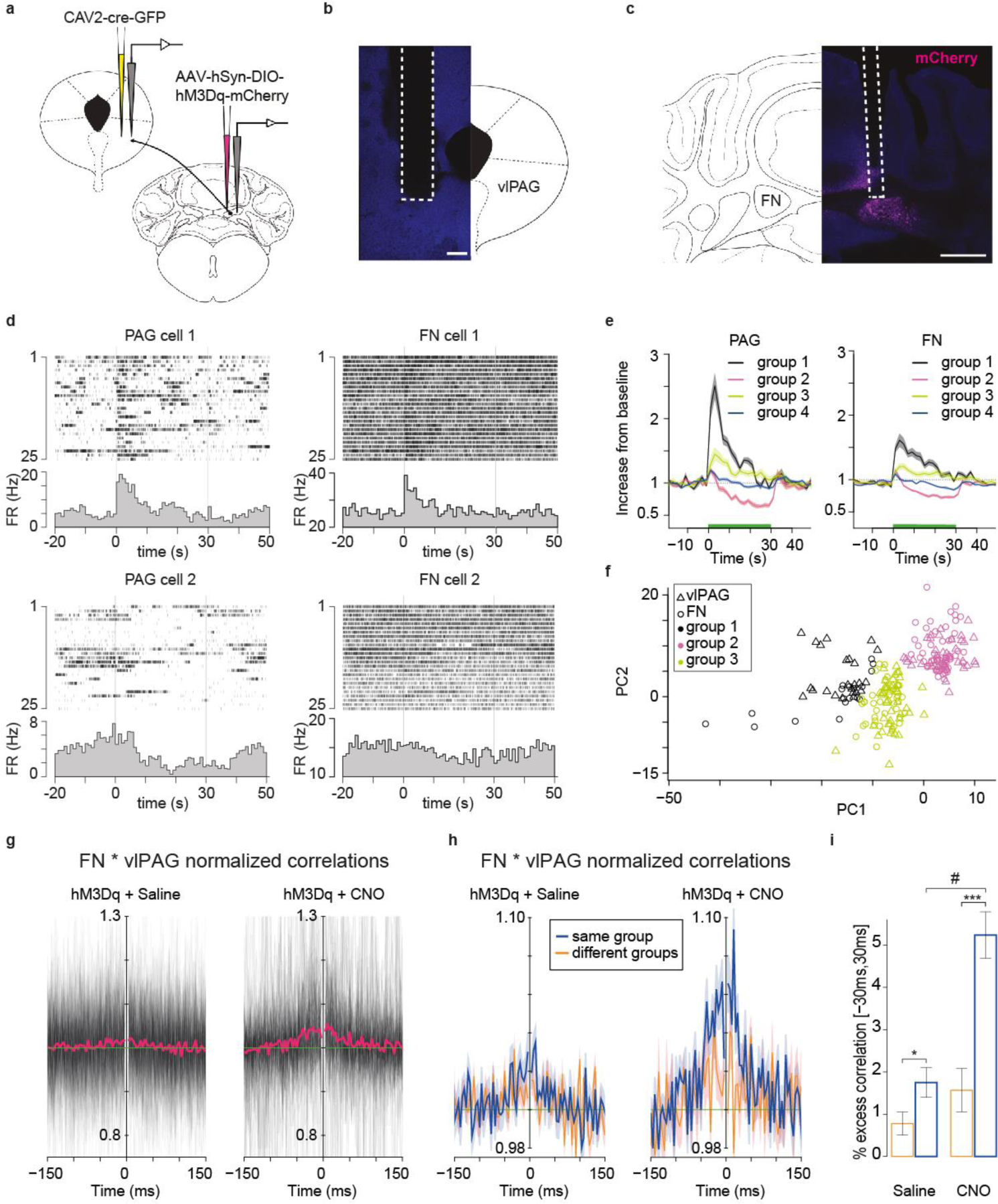
**a**, Cre-dependent expression of the hM3Dq in the FN-vlPAG projecting neurons and electrophysiological recording in FN and vlPAG in freely moving mice. **b,** Example of electrode position in the vlPAG (scale bar, 200 µm). **c,** Example of hM3Dq expression in FN-vlPAG projecting neurons and electrode position (scale bar, 500 µm). **d,** example rasters and peri-stimulus time histogram PSTH (grayed histogram) of the vlPAG and FN cells (vlPAG and FN cells 1 and cells 2 were each recorded from a same mouse). **e,** average PSTH profile of each group expressed as a fraction of the baseline. **f,** representation of the projection of each PSTH (for all days of recordings) on the two first principal component of the vlPAG PSTHs. The group 4 (non-modulated cells) is not represented on the plot. **g,** Superimposition of the PSTHs from all the pairs of FN-vlPAG units on the first day of extinction. The average is plotted in red. **h,** average of all normalized crosscorrelograms from pairs of FN-vlPAG units belonging either to the same group (blue) or different groups (orange) on the first day of extinction. S.e.m is indicated by the shaded area. Comparison of the crosscorrelograms averaged in a window [-30ms, 30ms]: Saline (*): F(1,208)=4.1, p=0.04, CNO (***): F(1,111)=19., p<0.001, Saline vs CNO for same group (#):F(1,10)=9.94, p=0.011. **i,** quantification of average correlations for pairs of cell belonging to the same or different groups.

To test if FN and vlPAG neurons belong to a functionally coupled network, we examined the crosscorrelations between the cells of the two structures. Indeed, many units showed a central peak corresponding to short term correlations (+/-30 ms), hence indicating near-synchronous activation (Fig. 6 g-i). These correlations did not show clear asymmetry (which would reveal directional entrainment), but we found that pairs of cells in the FN and in the vlPAG which belonged to the same functional clusters showed stronger short-term correlations than pairs of cells from the two structures belonging to different clusters (Fig. 6 h), consistent with the existence of several parallel circuits operating independently and involving specific FN and vlPAG subpopulations.

Furthermore, to evaluate whether the state of the FN-vlPAG projections regulates the fear learning at the time of the CS-US association ^37, 38^, we optogenetically stimulated the FN during the CS-US presentation (from the last 3 s of the CS until 3 s post CS-US, Fig.7 a, b, d). We found that mice exposed to the light stimulation exhibited a significant enhanced extinction of the fear response during the extinction sessions compared to the control group (Fig. 7 e). Moreover, the light stimulation had not effect on the immobility state in an open field (Fig. 7 c) or on the freezing behavior during the fear acquisition phase (Fig. 7 e). These results suggest that the cerebellum exerts a control over the fear memory formation through the FN-vlPAG pathway during the association between the CS and US.

**Figure 7.**
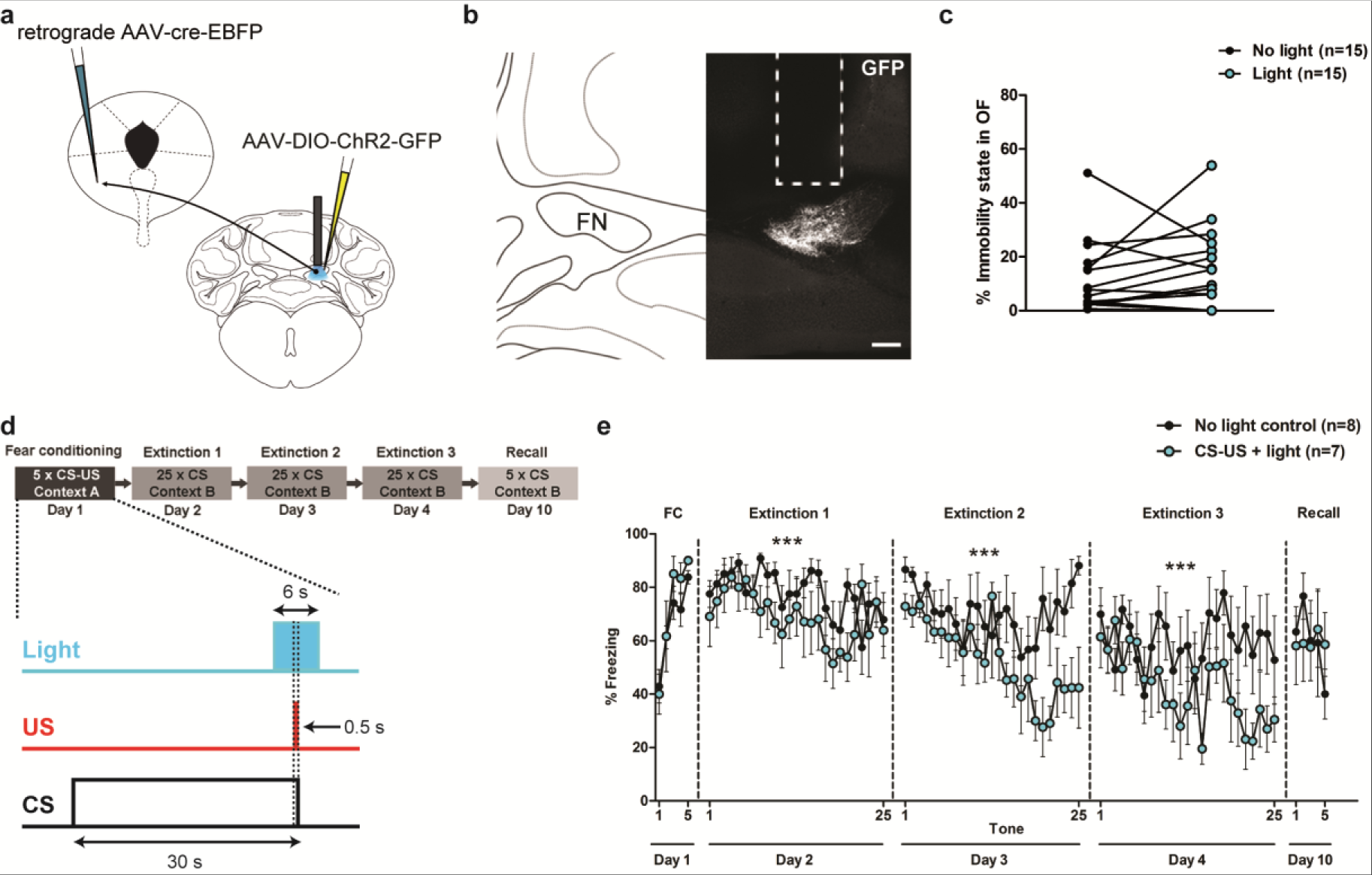
Involvement of FN-vlPAG pathway during the CS-US association. **a**, Strategy for optogenetic stimulation through an optic fiber in FN together with cre-dependent expression of ChR2-GFP in FN-vlPAG projecting neurons by expression of retrograde cre in vlPAG. **b,** Example of ChR2 expression in FN-vlPAG projecting neurons (scale bar, 200 µm). **c,** Light stimulation of FN-vlPAG projecting neurons have not effect on immobility state in the open field. Mice expressing ChR2 in FN-vlPAG projecting neurons were exposed to an open field 5 min without light and 5 min with light stimulation (n = 15, P > 0.05, paired t-test). **d,** Optogenetic stimulation protocol of FN-vlPAG projecting neurons during CS-US presentation in fear conditioning. Light was delivered every CS-US presentation, from the last 3 s of the CS to 3 s after the CS-US presentation. e, Freezing levels during fear conditioning and extinction sessions in control (no light stimulation, n = 8) and FN-vlPAG stimulated mice (CS-US + Light, n = 7). Mice that received light stimulation during CS-US pairing exhibited an enhanced extinction of fear response compared to the mice that were not stimulated (Extinction 1: F(1, 320) = 14.19, Extinction 2: F(1,292) = 38.3, Extinction 3: F(1, 317) = 1.06, P < 0.001, two-way ANOVA).

### Cerebellar input to the vlPAG contributes to extinction learning

Another interesting question is that if the cerebellum might also play a role in the extinction memory formation or in the retrieval of the fear memory. Therefore, in order to evaluate the extinction memory formation, mice expressing hM3Dq or hM4Di in FN were exposed to CNO during the first and second extinction sessions (Fig. 5 a-d). Mice fear conditioned were subjected to the first recall and extinction training the following day under CNO effect. During the first CS presentations, the different groups of mice exhibited the same levels of freezing, suggesting that CNO does not affect the fear memory retrieval (Fig. 5 g). Mice under FN-vlPAG activation did not exhibit differences in the freezing levels compared to the control group during the first extinction session. However, mice under inhibition of the FN-vlPAG projections exhibited impairment in the extinction of the fear response during the first and second extinction training (Fig. 5 g). Moreover, at the third extinction session, without exposure to CNO, these mice still exhibited higher levels of freezing compared to the control group, suggesting that FN-vlPAG projections could also control the extinction learning.

Taken together, these results indicate that FN inputs to the vlPAG are able to control in a bidirectional way the strength of the fear memories, reinforcing the association between the CS-US when this pathway is inhibited or diminishing CS-US association when it is activated. In addition, the inhibition of FN-vlPAG inputs could also impair the extinction learning.

### Inhibition of cerebellar input to PF reduce fear expression but not fear learning

We also evaluated the contribution of FN-PF pathway on fear learning, by analyzing the effect of activating or inhibiting this specific pathway during fear conditioning in mice expressing cre-dependent anterograde AAV-hM3Dq or -hH4MD injected in FN neurons, and retrograde CAV2-cre injected in the PF (Fig. 8 a-c). The inhibition of FN-PF projections induced significant increase in freezing behavior during the last CS-US presentation of the fear conditioning (Fig. 8 d), while activation of FN-PF pathway had not effect. Moreover, either activation or inhibition had not effect on the basal freezing levels in the context A before fear conditioning (Fig. 8 e). Nevertheless, mice that underwent FN-PF inhibition during fear conditioning expressed similar freezing levels during the extinction sessions compared to the control mice. Therefore, these results suggest that the FN-PF pathway might have a mild contribution on fear expression, but not on fear memory formation.

**Figure 8.**
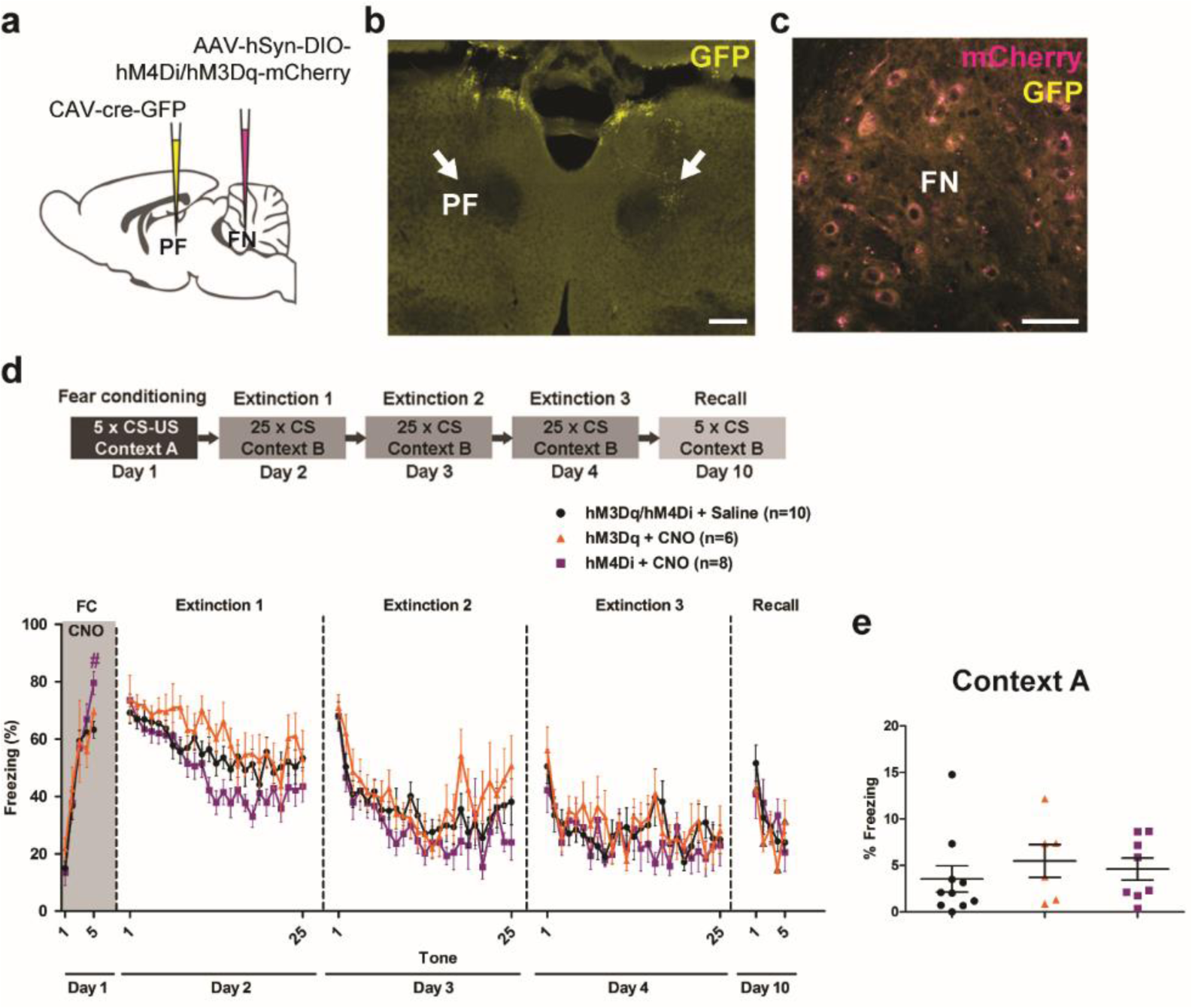
FN-PF pathway contributes on fear expression but not on fear memory formation. **a**, Chemogenetic modulation of FN-PF projecting neurons by bilateral cre-dependent expression of excitatory or inhibitory DREADDs (hM3Dq or hM4Di, respectively) in the FN in combination of CAV2-cre-GFP injection in the PF. **b,** Injection site of CAV2-cre-GFP in PF (arrows, scale bar, 200 µm). **c,** Cre-dependent expression of DREADDs in the FN (scale bar, 50 µm). **d,** FN-PF pathway stimulation (hM3Dq + CNO, n = 6) or inhibition (hM4Di + CNO, n = 8) during fear conditioning. Inhibition of FN-PF pathway induced an increase of freezing levels at the last CS presentation of the fear conditioning compared to the control group (hM3Dq/ hM4Di + Saline, n = 7; P < 0.05, Bonferroni post-hoc test). Inhibition or stimulation during fear conditioning had not effect on freezing behavior during the extinction sessions (P > 0.05, two-way ANOVA). Lines represent means ± s.e.m. **e,** Percentage of freezing in Context A before fear conditioning. Mice under inhibition or activation of the FN-vlPAG pathway exhibit similar basal levels of freezing to the context A compared to the control group (P > 0.05, one-way ANOVA). Scatter dot plot with mean ± s.e.m.

### FN-PF but not FN-vlPAG projections contributes to anxiety-like behavior

The neuronal circuits for fear and anxiety are overlapping and closely related ^7, 39^. Therefore, we evaluated if the FN-vlPAG or the FN-PF pathways contribute to anxiety behavior. We performed three different anxiety tasks: open field, elevated plus maze and dark-light box, to examine anxiety levels in mice under activation or inhibition of FN-vlPAG or FN-PF projections. There were not differences in anxiety parameters that indicate generalized anxiety-like behavior when FN-vlPAG pathway was activated or inhibited (Supplementary Tables 6 a). On the other hand, mice under activation of FN-PF pathway exhibited a decrease in the frequency on entries in the open arms in the EPM, and decreased frequency in the light zone in the light dark box test, indicating an anxiogenic effect induced by the activation of this pathway (Supplementary Tables 6 b). These results suggest that FN-PF projections modulate anxiety levels, while FN-vlPAG does not affect anxiety but it has a specific role on fear conditioning.

### FN-vlPAG and FN-PF pathways are not involved in painful sensitivity

PAG is involved in processing pain information from the periphery ^13^. Due to the activation or inhibition of FN-vlPAG pathway during CS-US presentations diminished or reinforced fear memory, respectively, we wondered if that could be the consequence of an alteration on the sensitivity to painful stimuli. Therefore, in order to evaluate the sensitivity of the mice under activation or inhibition of this pathway, we performed hot plate and tail immersion tests. No differences were found in the hot plate or the tail immersion sensitivities (Supplementary Fig. 6 c, d), indicating that FN-vlPAG inputs do not modulate the sensitivity to painful stimuli. Therefore, alteration in sensitivity is not responsible for the effect of FN-vlPAG inputs on fear memories formation and expression.

On the other hand, because FN-PF inhibition induced an increase in freezing expression during fear conditioning, it could be related to enhanced pain perception. However, no significant differences were found in sensitivity latencies in hot plate and tail immersion tests (Supplementary Fig. 6 e, f).

## Discussion

Despite many clinical and anatomical evidence indicating an involvement of the cerebellum in emotional functions ^19^, the nature of its contribution remains unclear. Here, we describe a link between the cerebellum and one element of the fear circuitry, the vlPAG, through which it exerts a bi-directional control over fear learning during the association of a cue and aversive events. Our experiments reveal that glutamatergic neurons in the FN send projections to the vlPAG where they form synapses onto glutamatergic and GABAergic neurons. The chemogenetic or optogenetic activation of this pathway during fear conditioning increases the activity of vlPAG neurons but yields reduced fear expression during the recall and extinction. In contrast, the inhibition of this pathway during fear conditioning results in a higher expression of fear and a slower extinction. Thus, our experiments are consistent with a participation of the cerebellum to the function of the vlPAG in fear learning.

The PAG has long been implicated as the organizer of behavioral components of the response to threat. The most characterized function of the vlPAG is its control of freezing via glutamatergic projections to pre-motor targets in the magnocellular nucleus of the medulla; this pathway is recruited in innate freezing, and in cued- or context-driven freezing respectively by GABAergic afferents from CEA ^10^ and glutamatergic afferents from the mPFC ^6^. The cerebellum may also participate to freezing expression. Indeed, the vlPAG has been shown to entrain (via an unknown pathway) the climbing fiber in the lobule 8 of the rat cerebellum, a region found to reduce both innate and cued freezing expression if lesioned ^24^. Such action on freezing expression is unlikely supported by cerebellum-vlPAG projections in our experiments: the chemogenetic or optogenetic stimulation of the FN-vlPAG did not affect the freezing evoked by the aversive stimulus during the presentation of the US, and failed to modify the cued-freezing at the time of recall (i.e. beginning of the extinction process). This makes unlikely that the FN-vlPAG pathway directly provides a strong entrainment to vlPAG neurons projecting to the magnocellular medulla-projecting, or to their afferent inhibitory neurons. Thus, the effects revealed in our work would not result from the regulation of the motor control of freezing.

The changes in freezing behavior observed in the three days following fear conditioning shall indeed result from a change in the learning of the CS-US association. A trivial explanation could be that the FN-vlPAG pathway modulates the sensory perception of the US at the time of fear encoding. Indeed, excitatory and inhibitory neurons in the vlPAG exert an anti-nociceptive effect ^10, 11^. However, we found that the same chemogenetic manipulations which alter the fear learning, failed to change the responses to pain in the tail-immersion and hot-plate tests, indicating that the changes in fear learning were not due to an effect of the FN-vlPAG directly on the antinociceptive circuit of the vlPAG. An appealing alternative possibility to explain the changes in fear learning observed during the extinction is an alteration of fear consolidation ^27^; however, the inactivation of the FN-vlPAG pathway after but not during conditioning failed to alter the retrieval of the fear response in the following days. Moreover, the optogenetic stimulation of the pathway only at the time of the CS-US presentation was sufficient to induce a decrease in the fear expression in the following days, consistent with a modulation on the encoding of the CS-US association and a limited contribution of the FN-vlPAG pathway to the consolidation process.

Increasing evidence points toward a bidirectional contribution of vlPAG to learning: the chemogenetic activation during learning subsequently reduces the cued feeding suppression indicating a reduced cued fear ^40^ (consistent with the chemogenetic and optogenetic findings of the present work), while the pharmacological inhibition of vlPAG before CS-US presentation may prevent learning of freezing responses ^41^. Some of the effect of the vlPAG on fear learning may involve a cued-antinociception triggered by an amygdala-vlPAG pathway ^12^, although we fail to observe a change in sensitivity to aversive stimuli in our chemogenetic experiments. However, the optogenetic inhibition of vlPAG has been found to inhibit the acquisition of conditioned fear response not only using aversive shocks but also using threatening visual stimuli ^37^, thus, pointing to a broad role of the vlPAG in fear learning. Indeed, a number of electrophysiological recordings and optogenetic manipulations indicate that vlPAG neurons encode positive fear prediction errors which determine the intensity of fear learning ^38, 41–43^.

The different types of vlPAG responses to the CS that we observed are consistent with previous reports: the fast onset neurons ^10, 12, 36, 44^ (our cluster 1) likely encode the fear prediction error ^42, 44^ and shall correspond, at least in part, to dopamine-vGluT2 expressing neurons ^42^. Slower excitatory responses (our cluster 3) rather encode threat probability ^12, 44^, while neurons with inhibitory responses to the CS (our cluster 2) more likely encode fear expression ^10, 45^.

A number of neurons in the FN exhibit responses to the CS homologous to the vlPAG responses, and pairs of neurons in the two structures with similar CS responses exhibit short-term (∼30ms) synchrony; the lack of sequential activation between the FN and vlPAG suggest that these activities result from the activation of larger network, possibly involving the amygdala. Indeed, the amygdala and the vlPAG have reciprocal connections that are essential for fear prediction error treatment ^12, 42^. However, the amygdala also projects heavily to the ponto-cerebellar system and is essential for cerebellar CS activity in reflex conditioning^25^, suggesting that fear prediction could be provided by the amygdala both to the vlPAG and cerebellum. Indeed, a recent imaging study has shown in a cued fear learning experiment that large portions of the cerebellar cortex are activated by fear prediction error^46^. The nature of cerebellar computations in fear prediction is still unknown, but the cerebellum provides an extensive representation of the context help to the massive mossy fiber system and fine temporal learning which may help to tune the fear prediction. The participation of the cerebellum to shock prediction is illustrated in aversive eyeblink conditioning where climbing fibers respond to the onset of the CS after learning ^47^.

Our study indicates a modulatory role of the FN-vlPAG pathway in fear extinction. Consistent with our results, evidence for negative prediction error was found in vlPAG ^40^, as well as in the ventral tegmental area and dorsal raphe ^48, 49^, which also receive inputs from the vlPAG.

A surprising finding of our work was the limited influence on fear learning of the most direct pathway between FN and amygdala via the PF. However, FN-PF activation induced anxiogenic behavior, which indicates that FN-PF could be involved in the modulation of anxiety states and could be related to the reported contribution of PF to observational social fear ^50^.

Understanding how cerebellar circuits engage with processing and associative learning of emotions in other areas of the central nervous system is an important avenue for future research ^17, 25^. Indeed, the cerebellar involvement in reward processing is engaging a pathway between the cerebellum and the ventral tegmental area ^17^. Our work shows that another pathway linking the cerebellum to vlPAG is controlling fear learning. The study of this pathway could be particularly relevant for the deeper understanding of emotional control and maladaptive threat processing.

## Materials and Methods Animals

Experimental subjects were adult C57BL/6N male mice, 8 to 12 week-old, wild-type (Charles River Laboratories) or mutant mice *Vglut2-cre* and *Glyt2-GFP* from an in-house colony (IBENS, Paris, France).

All mice were housed in groups of 4 mice per cage, in a 12 h light/dark cycle and all the experiments were performed in awake freely moving mice during the light cycle. Food and water were available *ad libitum*. All animal procedures were performed in accordance with the recommendations contained in the European Community Council Directives (authorization number APAFIS#1334-2015070818367911 v3).

### Surgeries and stereotaxic injections

Male mice 7 - 8 weeks old received a dose of buprenorphine and 15 min later they were deeply anaesthetized with isoflurane 3% and placed in a stereotaxic frame (Kopf Instruments). Anesthesia was kept constant with 1.5 - 2% isoflurane supplied via an anesthesia nosepiece, and body temperature maintained at 36°C with a heating pad controlled by rectal thermometer. A local analgesia with 1ml of 0.02% of lidocaine was injected subcutaneously above the skull, and then the skull was exposed and perforated with a stereotaxic drill at the desired coordinates relative to Bregma. For viral delivery, a capillary was lowered to coordinates just above the target area and 100-250 nl of virus solution was infused. After infusion, capillary was kept at the injection site for 5 more minutes before being slowly withdrawn. For viral injections and implants, the stereotaxic coordinates used were: vlPAG: AP: −4.45; ML: ± 0.55; DV: −2. −2.40; FN: AP: −6.37, ML: ± .7, DV: −2.12; BLA: AP: −1.35; ML: ±3.43; DV: −4.0; depths were taken relative to the dura.

### Neuroanatomical tracing

For neuroanatomical tracing mice were injected with 100 nl of anterograde AAV1-CB7-Cl-mCherry-WPRE-rBG (Upenn Vector) in the FN (n=14), 100 nl CTB-alexa 488 or -alexa 555 (Invitrogen) in the BLA (n=6), and 250nl CTB-CF594 (Biotium) or retrograde AAV-Syn-eGFP (Addgene) were injected in the vlPAG (n=6). For the CTBs infusions, mice were perfused the fourth day after the injections. For the AAVs infusions, mice were perfused 21 days after the injections. For synaptophysin labeling, 200 nl of AAV8.2-hEF1a-DIO-synaptophysin-GFP (Massachusetts General Hospital) was injected in FN of *vglut2-cre* mice (n=7), and a group of these mice were injected with 300 nl of AAV1.CAG.Flex.tdTomato.WPRE.bGH (Upenn Vector) in vlPAG (n=4), while the rest were used for gad67 immunostaining.

Mice were anesthetized with ketamine 80 mg/kg and xylazine 10 mg/kg, i.p., and the perfusions were performed with formalin (Sigma). Dissected brains were kept in formalin solution overnight at 4°C, and the brains were then stored in PBS solution at 4°C. Brains were sliced entirely at 90 µm using a vibratome (Leica VT 1000S), and mounted on gelatin-coated slides, dried and then coverslipped with Mowiol (Sigma). For counterstaining, sections were coverslipped with a mounting medium with DAPI (Fluoroshield, Sigma). Slices were analyzed and imaged using a confocal microscope (Leica TCS Sp8). In the same way, the placement of the optic fiber, electrodes and injections sites for all the experiments were assessed when the experiment was completed. Mice with no viral expression or misplacement of the optic fiber or electrode were excluded from the analysis.

### Immunohistochemistry

For the identification of GABAergic neurons in the vlPAG, mice were perfused with PFA 4%, stored overnight in PFA 4% at 4°C, and then stored in 25% sucrose solution in PBS. Brains were sliced coronally at 50 µm using a vibratome and slices were stored at −20°C in a cryoprotective solution until the moment of the immunostaining.

In order to analyze neuronal activity, immunostaining of the immediate early gene c-Fos was performed. Mice underwent fear conditioning protocol and 50 min after the end of the protocol each mice was perfused as has been described above.

For the immunostaining, selected slices were washed with PBS-0.3% Triton X-100, blockage of nonspecific sites was assessed by 2 h of incubation with 3% normal donkey serum (NDS) for GAD67-immunostaining (n=3) or with normal goat serum (NGS) for c-Fos-immunostaining (n=10). Sections were then incubated in a solution containing mouse anti-GAD67 (1:500, Milipore #MAB5406) in PBS-0.3% Triton X-100 with 1.5% NDS at 4°C for 72 h; or rabbit polyclonal anti-c-fos (1:800, Milipore #ABE457) in PBS-0.3% Triton X-100 with 1.5% NGS at 4°C for 24 h. After first antibody incubation, slices were rinsed and sections were 2 h incubated at room temperature with secondary antibody, donkey anti-mouse IgG conjugated to Alexa Fluor 488 or to Alexa 555 (1:400, Invitrogen), or goat anti-rabbit IgG-FITC (1:300, Jackson Immunoresearch). Slices were mounted on gelatin-coated slides with Mowiol and analyzed with the confocal microscope.

### Chemogenetics

To specifically modulate the activity of FN neurons projecting to vlPAG, excitatory or inhibitory DREADDs were expressed by bilaterally injection (200nl/injection site) of pAAV-hSyn-DIO-hM3Dq-mCherry (cre-dependent expression of excitatory DREADD, Addgene) or pAAV-hSyn-DIO-hM4Di-mCherry (cre-dependent expression of inhibitory DREADD, Addgene) in the FN in combination of 300nl CAV2-cre-GFP (Plateforme de Vectorologie de Montpellier) viral infusion in vlPAG. Surgeries and injections were performed 2 weeks before mice performed the behavioral tests. For neuronal modulation of animals expressing DREADDs, Clozapine N-oxide (Tocris Bioscience, 1.25mg/kg) was diluted in saline and injected i.p. 30 min before the start of the experimental session. Control group was injected with saline. Animals that received this treatment in experiments were habituated by saline injections during handling sessions.

### Pavlovian fear conditioning and extinction

Pavlovian fear conditioning was conducted in a context A: 17 × 17 × 25 cm chamber with black/white-checkered walls and a grid floor, with peppermint-soup odor, inside of a sound-attenuating chamber (Ugo Basile). The chamber was cleaned with a 70 % ethanol within mice. Following a 180 s acclimation period, mice received 5 pairings (90–120 s variable inter-pairing interval) between a 30 s, 80 dB, 2.7 kHz tone (conditioned stimulus, CS) and a 0.5 s, 0.4 mA scrambled footshock (unconditioned stimulus, US), in which the US was presented during the last 0.5 s of the CS. There was a 120 s no-stimulus consolidation period following the final CS-US pairing before mice were returned to the home cage. Stimulus presentation was controlled by the Ethovision XT 14 (Noldus).

Twenty-four hours later, recall of the response to the CS and the first extinction training was performed. Mice were placed in a novel context B: chamber with yellow semi-transparent cylindrical wall, a solid-Plexiglas, opaque floor, and vanilla-extract solution to provide a distinctive olfactory cue. The chamber was cleaned with the disinfectant detergent solution Surfa’safe premium (Anios) between different run. Following an initial 180 s acclimation period, the mouse received 25 × 30 s presentations of the CS (30 s no-stimulus interval). The second and third extinction sessions took place in the next two days under the same conditions. Finally, on day 10, recall test was performed in context B, by 5 × 30 s presentations of the CS (30 s no-stimulus interval).

Mice were videotracked using Ethovision XT 14 during all the trial. Freezing (no visible movement except that required for respiration) was assessed by the inactive periods, defined as periods of time during which the average pixel change of the entire video image was less than 0.5 % (from one video frame to another one) and the threshold was fixed avoiding the detection of the breathing movements. Freezing behavior was analyzed during each CS presentation and during the habituation period to the context A and B.

### Anxiety tests

#### Open field test

To evaluate anxiety-like behavior and locomotor activity, mice were placed in a 38 cm diameter circular arena (Noldus) and recorded by video from above. The position of the center of gravity of the mice was tracked with Ethovision XT 14 and the distance moved, time in the center area, frequency of entries in the center area, were analyzed for a 20 min period.

#### Elevated plus maze

Mice were placed in the center zone (6 x 6 cm), facing an open arm of an elevated plus maze (Noldus, elevated 52 cm above the floor) with 2 open arms (36 cm length, 6 cm width) and 2 wall-enclosed arms (closed arms, 36 cm length, 6 cm width, walls 25 cm high) and let explore freely for 5 minutes. Their path was videotracked using Ethovision XT 14, and the amount of time spent and distance moved in the open arms, closed arms and center zone were evaluated.

#### Light-dark box

Mice were placed in the light zone and let explore the light-dark arena (Noldus) freely for 5 minutes. Their path was videotracked in the light zone using Ethovision XT 14 and the amount of time spent in the light versus dark zones and distance travelled in the light zone were evaluated, as well as the latency until they escaped to the dark zone. Lux levels were 150 lux in the light zone and about 0 lux in the dark zone. Light zone was 40 cm x 20 cm in size, and dark zone was 20 x 20 cm.

### Hot plate and tail immersion tests

Effect on sensitivity was tested one week after recall test using a hot plate analgesia meter (Harvard apparatus) and tail immersion test. Mice received a CNO or saline i.p. injection and 30 min after were put on the hot plate at 55°C. The experiment was stopped as soon as the mice performed the first licking or jump. Mice were videotracked and the latency to the first reaction (hindpaw shaking or licking, or jump) was recorded.

Tail immersion test was performed after hot plate test. Mice tale tips were immersed in hot water with a temperature of 50°C, and tail withdrawal latency was recorded.

### Optogenetics and electrophysiological recordings

For optical manipulation and electrophysiological recordings, mice were injected with 200 nl of cre-dependent AAV-ChR2-GFP (Addgene) in the FN combined with local injection of 200 nl of retrograde AAV-cre-EBFP (Addgene) into the vlPAG. After bilateral infusion, incisions were sutured closed using nylon monofilament. The cerebellar FN preferentially sends projections to the contra-lateral vlPAG regions (Fig. 1 a, b). Therefore, in order to compare responses we recorded neuronal activity in both contra- and ipsi-lateral vlPAGs when we unilaterally stimulated the FN. To record cell activity in the vlPAG we used bundle of electrodes consisting of nickel/Chrome wire (0.004 inches diameter, Coating 1/4 Hard PAC) folded and twisted into bundles of 6 electrodes. Pairs of bundles were inserted in metal cannulas (stainless steel, 29 G, 8 mm length, Phymep) in order to protect them and to guarantee a correct placement into the brain. Cannulas and bundles were also attached to an electrode interface board (EIB-16; Neuralynx, Bozeman) by Loctite universal glue in a configuration allowing us to record vlPAG. The microwires of each bundle were connected to the EIB with gold pins (Neuralynx). The entire EIB and its connections were fixed in place by dental cement (Super Bond). The impedance of each electrode was measured and the 1 kHz impedance was set to 200– kΩ using gold-plating (cyanure-free gold solution, Sifco). During the surgery, the electrode bundles were inserted into the brain and the ground was placed over the cerebellum, between the dura and the skull.

To perform optical stimulation, mice were implanted unilaterally with an optical fibre (0.22 numerical aperture, Thor labs) housed in a stainless steel ferula (Thor Labs) just dorsal to the FN (Supplementary fig. 3). Ferrules were adhered to the skull with adhesive, and then a protective head cap was constructed using dental cement. The viral injections, optical fiber and electrodes implantation were performed in the same surgery and the optical manipulation and recordings were performed 3 weeks after surgery to ensure ChR2 expression.

For the ramp of optical activations, electrophysiological recordings in the vlPAG were performed in freely moving mice placed in an open field. Recordings were performed using an acquisition system with 16 channels (sampling rate 25 kHz; Tucker Davis Technology System 3, Tucker-Davis Technologies) as described ^51, 52^. Mice received a stimulationat 0.5 Hz, 50 ms, and the light intensities used were 0.37, 0.54, 0.75, 1.03 and 1.33 mW (at the fiber tip). The signal was acquired by a headstage and amplifier from TDT (RZ2, RV2, Tucker-Davis Technologies). The spike sorting was performed with Matlab (Mathworks) scripts based on k-means clustering on PCA of the spike waveforms (Paz et al. 2006). At the end of the experiments, the position of the electrodes was verified by electrolytic lesions and comparison to a brain atlas after brain slicing was performed.

For the optogenetic activation of the FN during fear conditioning, mice were randomly designated to non-stimulated or stimulated group. The stimulated group received on each of the 5 CS-US presentations 20 ms light pulses at 10 Hz, 20 ms, 1 mW during a total of 6 s, from the last 3 s of the CS until 3 s posterior to the end of CS-US presentation (Fig. 7 d).

### Chemogenetics and electrophysiological recordings

Mice were injected bilaterally with 300nl CAV2-cre-GFP in the vlPAG, and with 200 nl of pAAV-hSyn-DIO-hM3Dq-mCherry or pAAV-hSyn-DIO-hM4Di-mCherry in the FN, as has been detailed above (see Chemogenetics section). After viral infusions, electrodes were placed in FN and vlPAG as was described above (see Optogenetics and electrophysiological recordings section). Fear conditioning was performed 2-3 weeks after the surgeries, and electrophysiological recordings during extinction sessions were analyzed.

### Statistical analysis

The effect of activation or inhibition of FN-vlPAG projections was analyzed using analysis of variance (ANOVA) and Newman-Keuls or Bonferroni post-hoc comparisons, or paired t-test, where appropriate. All statistics were performed in Graph Pad Prism® (Version 5) and RStudio®, unless otherwise indicated, and all statistical tests used are indicated in the figures legends. The effect of trial on freezing during fear conditioning and extinction was analyzed using two-way ANOVA. Experimental designs with two categorical independent variables were assumed to be normal and analyzed by two-way ANOVA without formally testing normality. All significance levels are given as two-sided and were corrected for multiple comparisons, wherever applicable. Statistical significance was set at *P* < 0.05.

To analyze the response to the optogenetic stimulations, we constructed the PSTH (bin 2ms) and converted these PSTH to a z-score using the pre-PSTH period (subtracting by baseline mean and dividing by the baseline standard deviation), the latency was determined by the time of the first bin with strongly significant z-score (>4).

To distinguish the different types of neuronal responses to the CS, we first identified the cells which were significantly modulated by the CS. For this purpose, we constructed the peri-stimulus time histogram (PSTH, 2 s bin), with a 20s baseline. The PSTH were then converted to z-score by subtracting the mean of the baseline and dividing by its standard deviation. Cells with less than one significant bins (z-score >3) were classified as weakly- or not-modulated. The remaining PSTH from vlPAG neurons were then splited in 3 groups by k-mean clustering; the number of clusters was determined using the R package NbClust ^53^with the Ward method (which minimizes the total within-cluster variance). The FN neurons were then classified by minimizing their distance to the PAG centroids. The FN-PAG crosscorrelograms were computed for the 5-min time period preceding the first CS, all the 30s time periods during all the CS, and the 30s time period after each CS. Each crosscorrelogram was normalized by the average of the crosscorrelogram values in the [−150ms,−100ms]U[100ms,150ms] interval. The excess correlations were computed as the average crosscorrelogram values in the [-30ms,30ms] interval. Statistical significance was tested by repeated-measure ANOVA (mice or cell pair nested in mice grouping factors as appropriate).

### Data availability

The data that support the findings of this study (Fig. 1-8, Supplementary Figs. 1-5) are available from the corresponding author upon reasonable request. For additional information please refer to the corresponding Life Sciences Reporting Summary.

## Code availability

The code for analysis is available from the corresponding author upon reasonable request.

## Acknowledgments

This work was supported by Fondation pour la Recherche Medicale (FRM, DPP20151033983) to D.P. and Agence Nationale de Recherche to D.P. (ANR-16-CE37-0003-02 Amedyst, Labex Memolife) and to C.L. (ANR-12-BSV4-0027, ANR-17-CE37-0009 Mopla) and by the Institut National de la Santé et de la Recherche Médicale (France). IAG was supported by the project 819/11.01.2019 by UMF Carol Davila, Bucharest. The authors declare no competing financial interests. We thank the Imaging Facility at IBENS.

## Author contributions

JLF and DP designed the experiments; JLF, CL and DP wrote the manuscript; JLF, RWS and CL analyzed the data; JLF, HBA and CMH performed the experiments, HBA, CMH, RWS and IAG provided technical support.

---

**Supplementary figure 1.**
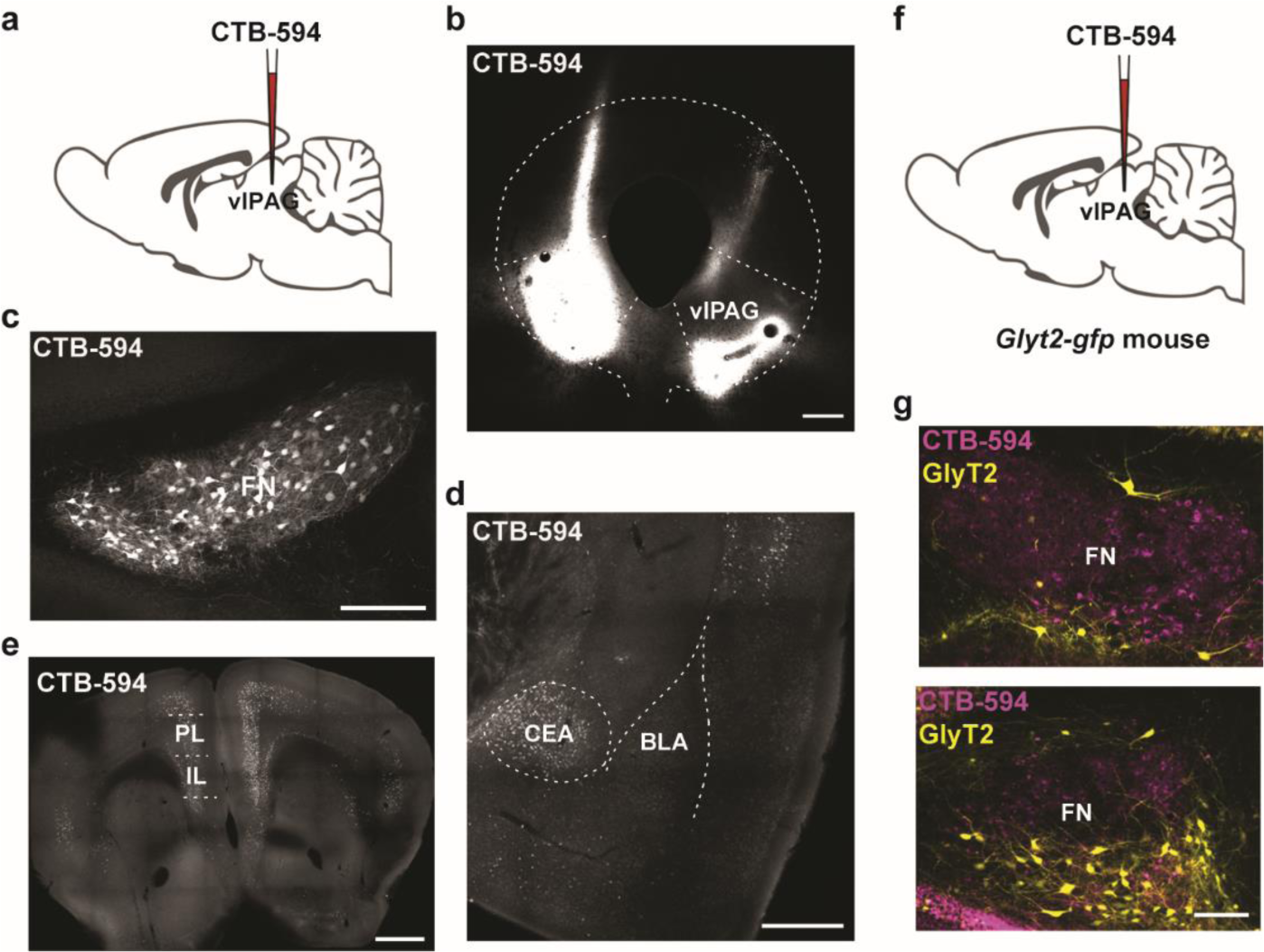
vlPAG receives monosynaptic projections from FN, CEA, and mPFC. **a**, Retrograde tracing from vlPAG. **b,** Injection site of CTB-594 in the vlPAG (scale bar, 100 µm). **c-e,** vlPAG received monosynaptic projections from FN (scale bar, 200 µm), CEA (scale bar, 500 µm), and from the PL and IL areas of the mPFC (scale bar, 1mm). **f,** Retrograde injection of CTB-594 in vlPAG of glyt2-gfp mice line. **g,** Retrograde tracing from vlPAG into the FN did not co-localized with GlyT2+ cells (scale bar, 100 µm).

**Supplementary figure 2.**
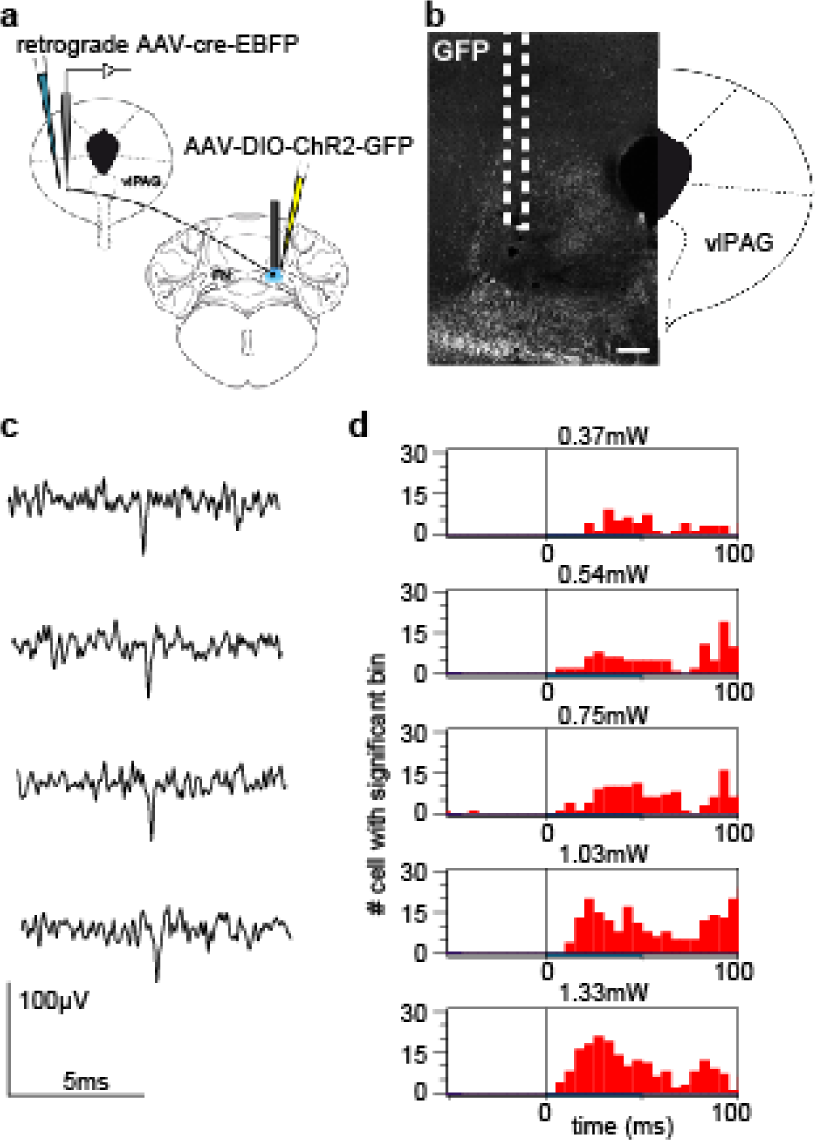
Optogenetic stimulation of FN-vlPAG pathway and electrophysiological recordings in freely moving animals. **a**, Strategy to assess optogenetic stimulation in FN-vlPAG projecting neurons and electrophysiological recordings in the vlPAG. **b,** Example of electrode position in the vlPAG for the electrophysiological recordings (scale bar, 200 µm). **c,** same trace excerpts as fig 3b, after high-pass filtering. **d,** histogram of number of cells exhibiting a significant bin (z-score PSTH > 4) at each time around the stimulation (blue bar under the histogram) for the range of

**Supplementary figure 3.**
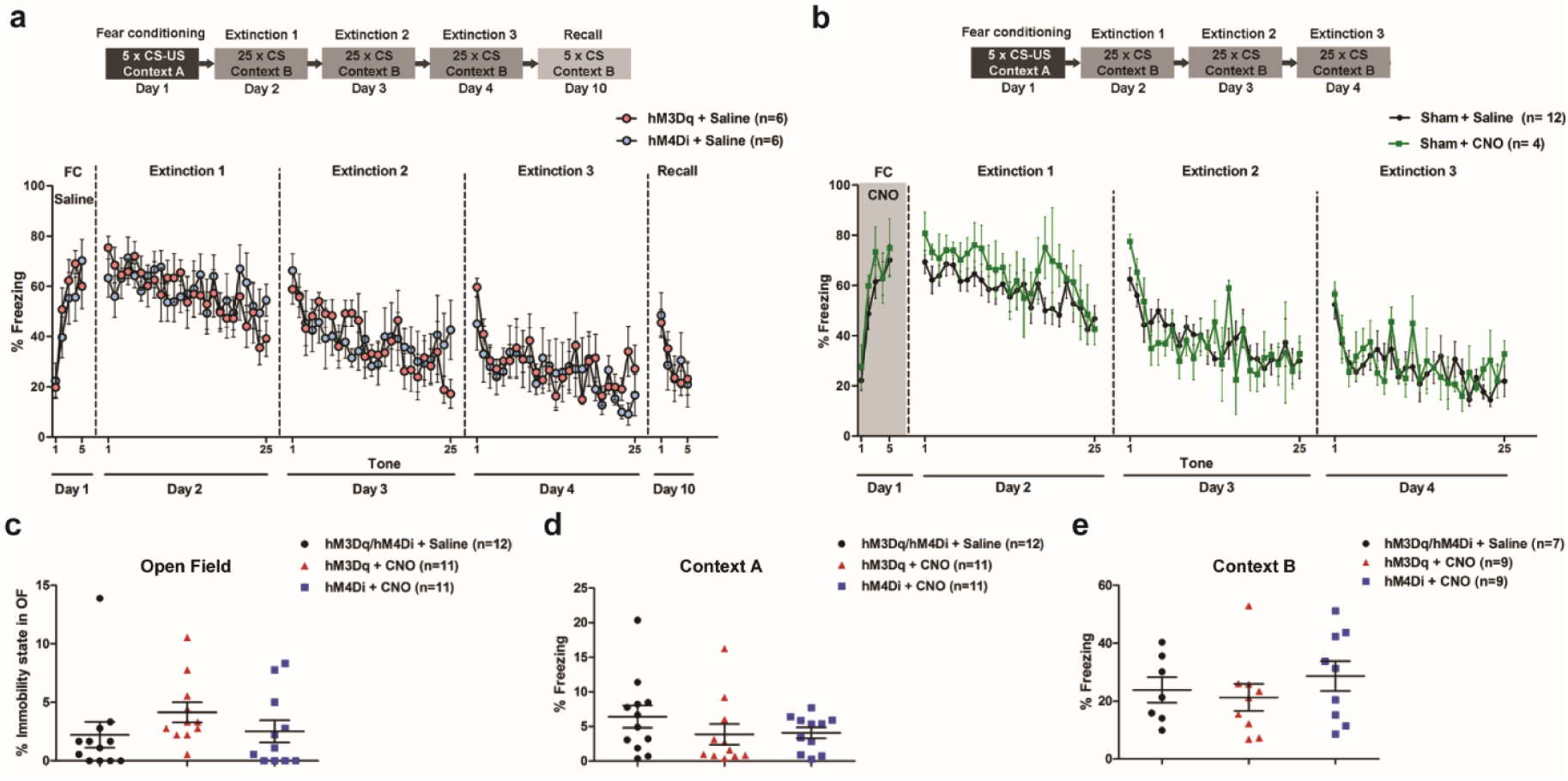
**a**, No differences were found in fear conditioning and extinction between control mice injected with saline expressing hM3Dq and hM4Di (hM3Dq + Saline, n = 6; hM4Di + saline, n = 6; P > 0.05, two-way ANOVA), lines represent means ± s.e.m. **b,** CNO had not effect *per se* on the freezing behavior in control sham mice (Sham + saline, n = 12; Sham + CNO, n = 4; P > 0.05, two-way ANOVA), lines represent means ± s.e.m. **c,** Chemogenetic activation or inhibition of FN-vlPAG pathway have not effect on immobility state in the open field (hM3Dq/ hM4Di + Saline, n = 12; hM3Dq + CNO, n = 11; hM4Di + CNO, n = 11; P > 0.05, one-way ANOVA). **d,** Mice under chemogenetic inhibition or activation of FN-vlPAG pathway exhibited similar basal freezing levels in Context A before fear conditioning (hM3Dq/ hM4Di + Saline, n = 12; hM3Dq + CNO, n = 11; hM4Di + CNO, n = 11; P > 0.05, one-way ANOVA), **e,** Mice under chemogenetic inhibition or activation of FN-vlPAG pathway during extinction exhibited similar basal freezing levels to the Context B before extinction training (hM3Dq/ hM4Di + Saline, n = 7; hM3Dq + CNO, n = 9; hM4Di + CNO, n = 9; P > 0.05, one-way ANOVA).

**Supplementary figure 4.**
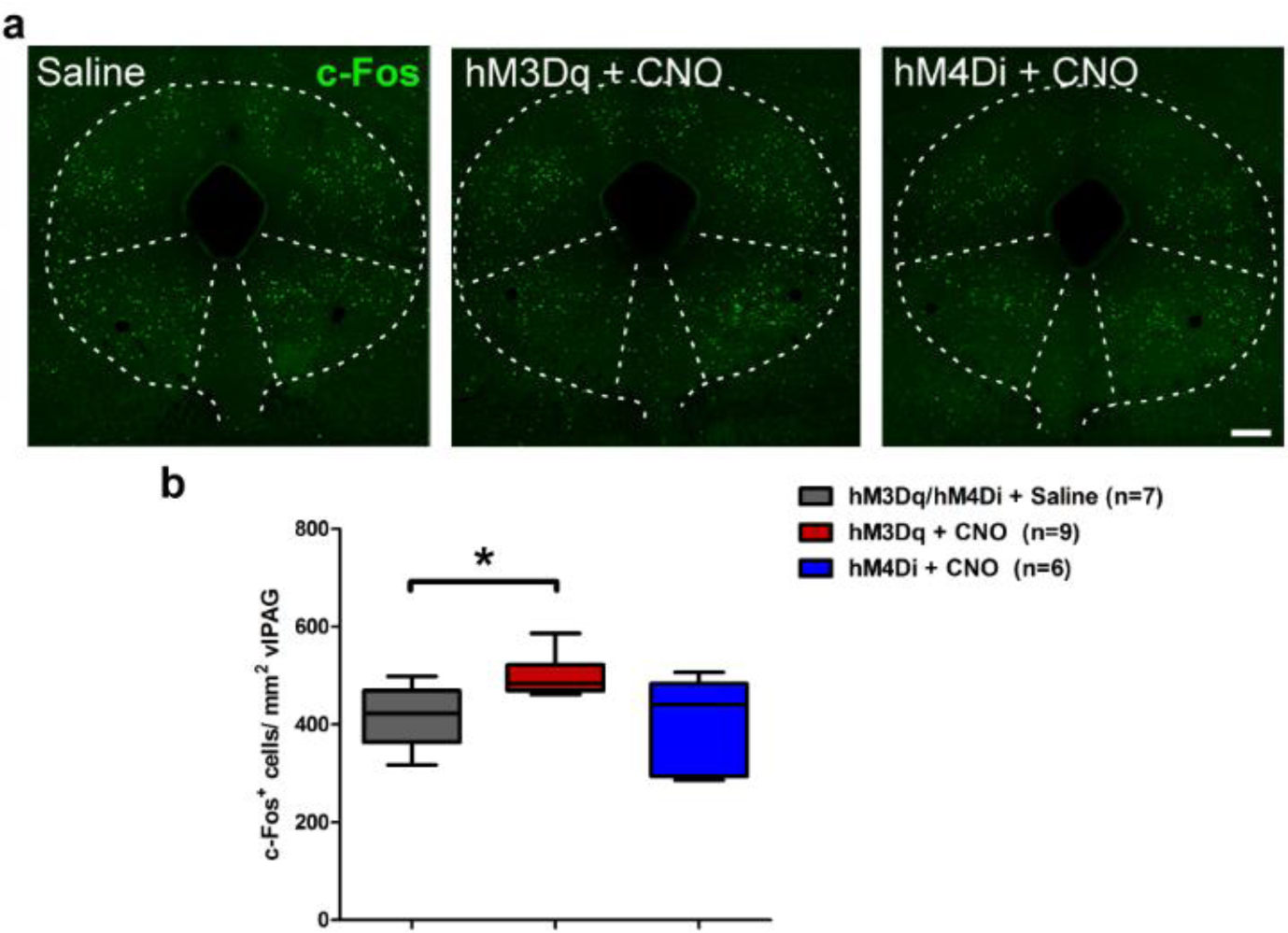
Stimulation of glutamatergic FN-vlPAG projections during fear conditioning increase neuronal activity in the vlPAG. **a**, C-Fos expression in vlPAG after fear conditioning under FN-vlPAG chemogenetic stimulation and inhibition (scale bar, 200 µm). **b,** C-Fos expression increased in vlPAG under stimulation of FN-vlPAG pathway during fear conditioning (hM3Dq/ hM4Di + Saline, n = 7; hM3Dq + CNO, n = 9; hM4Di + CNO, n = 6). Box represents quartiles and whiskers min to max. P < 0.05, Newman-Keuls Multiple comparison test).

**Supplementary figure 5.**
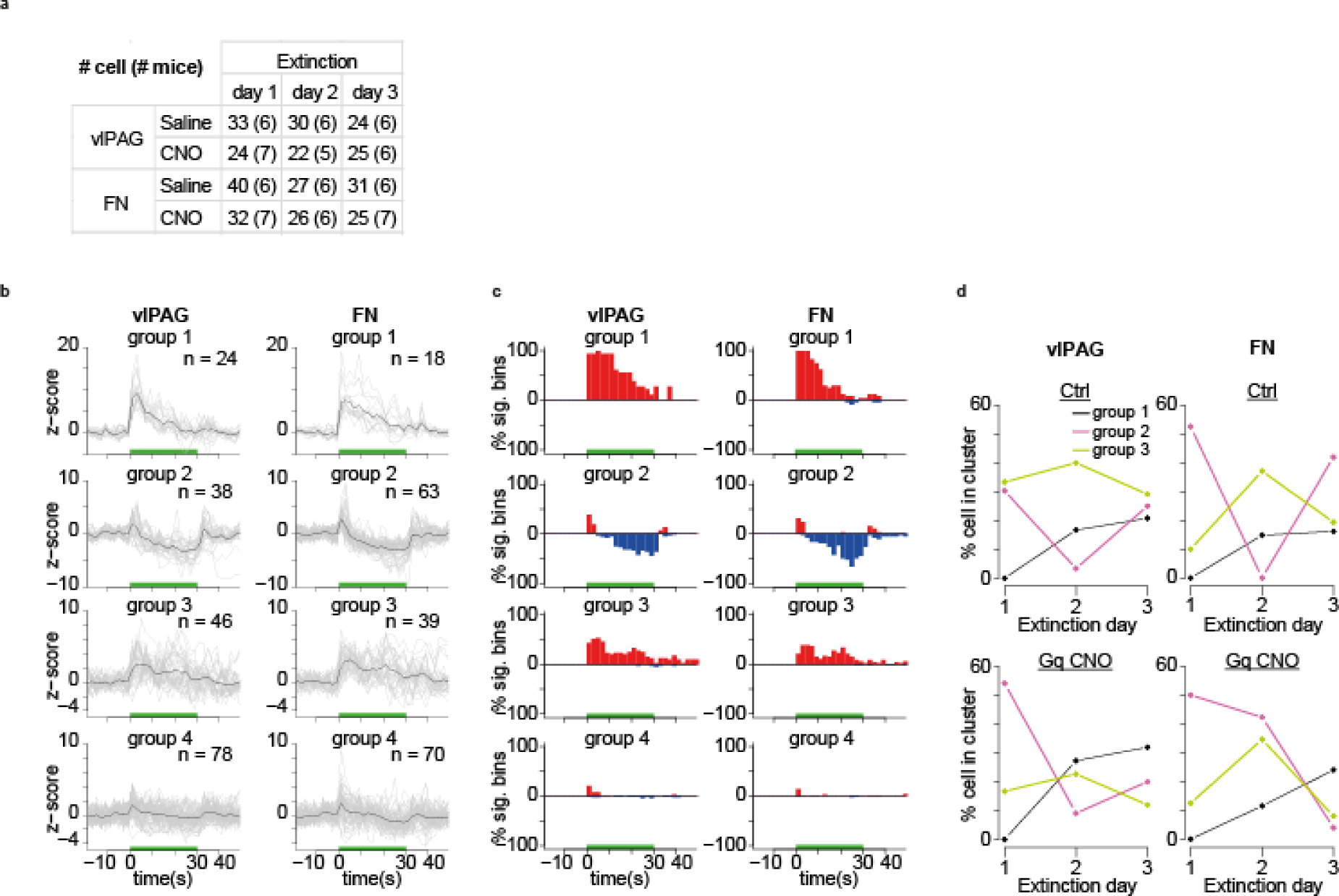
Summary of the firing poperties of the vlPAG and FN units recorded. **a,** count of cell and mice (number in parenthesis) across days. **b,** PSTH of all the units sorted by groups (n indicates the cell count). **c,** percentage of cell for each group with positive (red) and negative (blue) significant deviation from baseline for each time bin of the PSTH. **d,** distribution of the fraction of cells belonging to each group.

**Supplementary figure 6.**
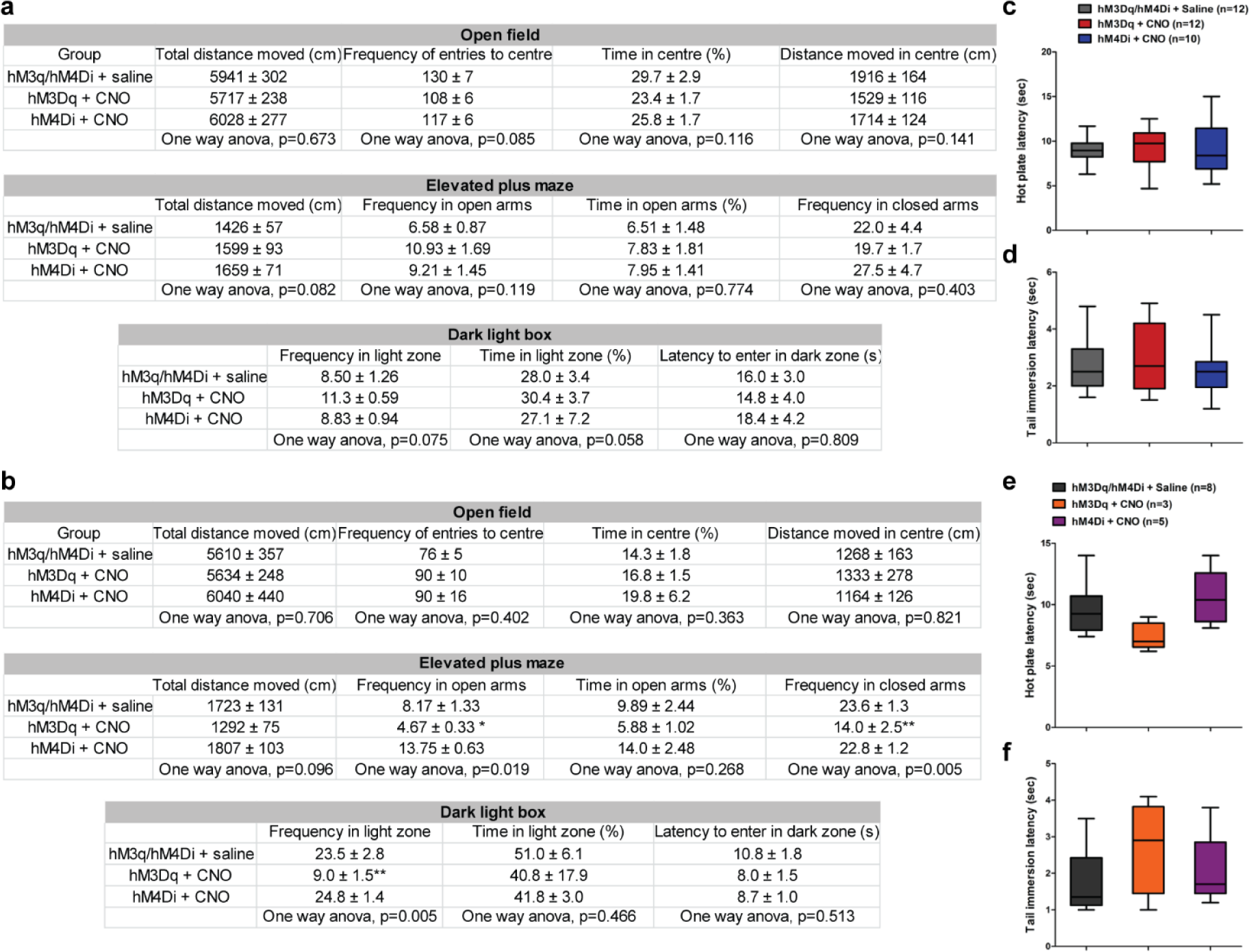
FN-PF, but not FN-vlPAG, is involved in anxiety like behavior, and both have not effect on pain sensitivity. **a**, FN-vlPAG chemogenetic activation or inhibition did not induce anxiety-like behavior (P > 0.05, one-way ANOVA), means ± s.e.m. **b,** FN-PF activation induced an increase in anxiogenic like behavior in the Elevated plus maze and in the Dark light box tests (P < 0.05, Bonferroni post-hoc test), means ± s.e.m. **c,d**, FN-vlPAG (hM3Dq/ hM4Di + Saline, n = 12; hM3Dq + CNO, n = 12; hM4Di + CNO, n = 10; P > 0.05, one-way ANOVA) and **e, f,** FN-PF (hM3Dq/ hM4Di + Saline, n = 8; hM3Dq + CNO, n = 3; hM4Di + CNO, n = 5; P > 0.05, one-way ANOVA) stimulation or inhibition had not significant effect on hot plate and tail immersion tests. Box represents quartiles and whiskers min to max.

